# Human NLRP3 inflammasome activation leads to formation of condensate at the microtubule organizing center

**DOI:** 10.1101/2024.09.12.612739

**Authors:** Jue Wang, Man Wu, Venkat G. Magupalli, Peter D. Dahlberg, Hao Wu, Grant J. Jensen

## Abstract

The NLRP3 inflammasome is a multi-protein molecular machine that mediates inflammatory responses in innate immunity. Its dysregulation has been linked to a large number of human diseases. Using cryogenic fluorescence-guided focused-ion-beam (cryo-FIB) milling and electron cryo-tomography (cryo-ET), we obtained 3-D images of the NLRP3 inflammasome *in situ* at various stages of its activation at macromolecular resolution. The cryo-tomograms unexpectedly reveal dense condensates of the human macrophage NLRP3 inflammasome that form within and around the microtubule organizing center (MTOC). We also find that following activation, the trans-Golgi network disperses and 50-nm NLRP3-associated vesicles appear which likely ferry NLRP3 to the MTOC. At later time points after activation, the electron-dense condensates progressively solidify and the cells undergo pyroptosis with widespread damaged mitochondria and autophagasomal structures.

## Introduction

Canonical inflammasomes are cytoplasmic supramolecular complexes responsible for detecting and sensing infections and cellular distress (1–3). Upon activation, these multi-protein complexes assemble into supramolecular structures that activate pro-caspases, release proinflammatory cytokines, and eventually cause pyroptotic cell death (1–3). One of the most extensively studied members in this family, the NLRP3 inflammasome, is composed of an NLRP3 sensor protein (also known as cryopyrin), an apoptosis-associated speck-like protein containing a C-terminal caspase recruitment domain (ASC) adaptor, and the pro-caspase-1 effector (1, 2, 4, 5). Early studies revealed that germline mutations and hyperactivity of NLRP3 are associated with rare hereditary autoinflammatory diseases in humans (6–8). NLRP3 has also now been implicated in the development of many common diseases including Parkinson’s disease, Alzheimer’s disease, cardiovascular diseases, and age-related metabolic complications (1, 6–9).

In macrophages, activation of the NLRP3 inflammasome involves two steps. In the first step (priming), Toll-like receptors (TLRs) on the cell surface recognize pathogen-associated molecular patterns (PAMPs) or damage-associated molecular patterns (DAMPs). This leads to the production of inflammasome proteins including NLRP3 and the precursor of proinflammatory cytokine IL-1β (1, 2, 4, 5). In the second step, bacterial toxins like nigericin, extracellular ATP, or other molecules induce cellular K^+^ efflux and NLRP3 activatiion. Active NLRP3 recruits ASC, which is thought to polymerize and thereby cause caspase-1 processing via proximity-induced dimerization and self-cleavage (1–3, 5, 10). Active caspase-1 cleaves the precursors of IL-1β and IL-18, leading to their maturation (11). Active caspase-1 also cleaves the pore-forming protein gasdermin D (GSDMD), which then forms pores in cytoplasmic membranes that release the processed cytokines and induce pyroptosis (12).

By fluorescence microscopy, the NLRP3 inflammasome is seen as a single speck within each cell (13–15). Some of us (Magupalli et al.) and others showed previously that this single speck contains the caspase-1 holoenzyme and colocalizes with the microtubule-organizing center (MTOC) (9, 14, 16). Many in-vitro structures of purified components and reconstituted sub-assemblies are already available, including cryo-EM structures of inactive, monomeric NLRP3 (in complex with NEK7) (17, 18); oligomeric NLRP3 (18–21); and NLRP3 in active-disc form (in complex with NEK7 and ASC) (10). A recent in situ cryo-ET study of ASC-overexpressing mouse immortalized bone-marrow-derived macrophages (iBMDMs) (22) revealed networks of ASC filaments (23), consistent with reports that purified ASC pyrin domain (PYD) and caspase-recruitment domain (CARD) assemble into filaments in vitro (24, 25). However, it remains unclear if they were an artifact of ASC over-expresssion.

Cryo-FIB milling and cryo-ET have become powerful techniques to visualize protein complexes in their near-native state to macromolecular resolution (26, 27). Previous studies have successfully targeted certain cells, organelles, and large protein aggregates by fluorescence (23, 27, 28). However, axial guidance of the milling remains challenging for small and rare structures, like the MTOC or the NLRP3 inflammasome, due to registration errors and motion of the sample that occur both during the transfer of the sample from the cryogenic light microscope to the FIB-SEM and during the milling process itself. These challenges persist even when the light microscope is integrated into the FIB-SEM vacuum chamber due to the common configuration of the optical focus and the coincident point of the FIB-SEM being in two distinct regions of the vacuum chamber. Here we empolyed a new, tri-coincident cryo-FIB-SEM-flourescence instrument (29) to overcome these challenges. Using real-time simultaneous fluorescence and cryo-FIB to guide the milling and subsequent cryo-ET, we visualized NLRP3 inflammasomes directly in situ at the MTOC in human macrophages at various stages of activation.

## Results

### Fluorescence-guided cryo-FIB milling

In order to find the NLRP3 inflammasome (hereafter referred to as simply the inflammasome) *in situ* and guide the FIB-milling, we first constructed a cell line with a fluorescent marker, mScarlet, fused to NLRP3 (**Fig. S1A**). This was done in human THP-1 monocytic cells in which endogenous NLRP3 had been knocked out. Monocytic cells were then differentiated into macrophages. This macrophage cell line recapitulated all the hallmark behaviors of wildtype inflammasome activation: after priming with lipopolysaccharides (LPS) that activate TLR4 and stimulating with nigericin (a potassium ionophore frequently used to activate NLRP3 (30, 31)), we observed ASC speck formation, GSDMD processing, and cell death as shown by LDH release (**Fig. S1B-D**). These effects were seen in WT and mSL-NLRP3 cells, but not in the parent NLRP3-knock-out cells, confirming the dependence of the processes on NLRP3 (**Fig. S1B-D**).

To prepare samples for cryo-ET we milled ∼200-nm-thick lamellae (29) within cells at various time points before and after priming and stimulation using the ENZEL tri-coinicident cryo-FM and cryo-FIB-SEM platform. This stage allows the milling process to be guided in realtime by fluorescence, and with multiple color channels (29). Thus, in addition to using mScarlet to mark the inflammasome, we also added SiR-tubulin to mark the MTOC (32). Untreated cells exhibited only SiR-tubulin signal. Primed cells exhibited both mScarlet (NLRP3) and SiR-tubulin signals. Addition of nigericin to primed cells caused the mScarlet fluorescence to condense into a single bright punctum colocalized with the MTOC (**Fig. S1E, Fig. S2**). aUsing this tri-coincident cryo-FIB-milling system, 52 of the 82 total lamella milled retained the target of interest, as seen by fluorescence of the final lamella and cryo-ET.

### NLRP3 accumulates into a condensate at the MTOC

SiR-tubulin fluorescence was seen to precisely correlate with centrioles in the cryo-tomograms, marking the MTOC unambiguously (**Fig. 1A-B**). In nigericin-stimulated cells, but not in untreated or only LPS-primed cells, an electron dense region containing irregular patch-like densities was seen between and around the centrioles, expanding approximately 2 µm in diameter, (**Fig. 1C-D**; **Fig. 2, H-J; Fig. S3A**). The patches were variable in size and shape, with rough dimensions of 30 to 200 nm (**Fig. S3B**). mScarlet fluorescence correlated with this electron dense region, identifying it as containing large amounts of NLRP3 (**Fig. 1C-D**; **Fig. 2C-E, H-J; Fig. S3A**). While the region around the MTOC contained ribosomes before stimulation, ribosomes were excluded afterwards. Thus the NLRP3 inflammasome punctum correlates to a patchy, electron-dense and ribosome-free condensate, similar to other cellular condensates (33, 34). As shown below, the numbers of small vesicles as well as microtubule nucleation sites also decreased with time after nigericin stimulation, another indication that the condensate gradually excluded other cellular structures (**Fig. 3A**). No extended filaments besides microtubules were observed, unlike an organized network of tubular filaments seen by cryo-ET under an ASC-overexpressing condition (23), At later time points after nigericin stimulation (> 30 min), fewer small patches were seen, replaced instead by large, more uniformly-dense condensate (**Fig. 1D**; **Fig. 2E**).

**Fig. 1.**
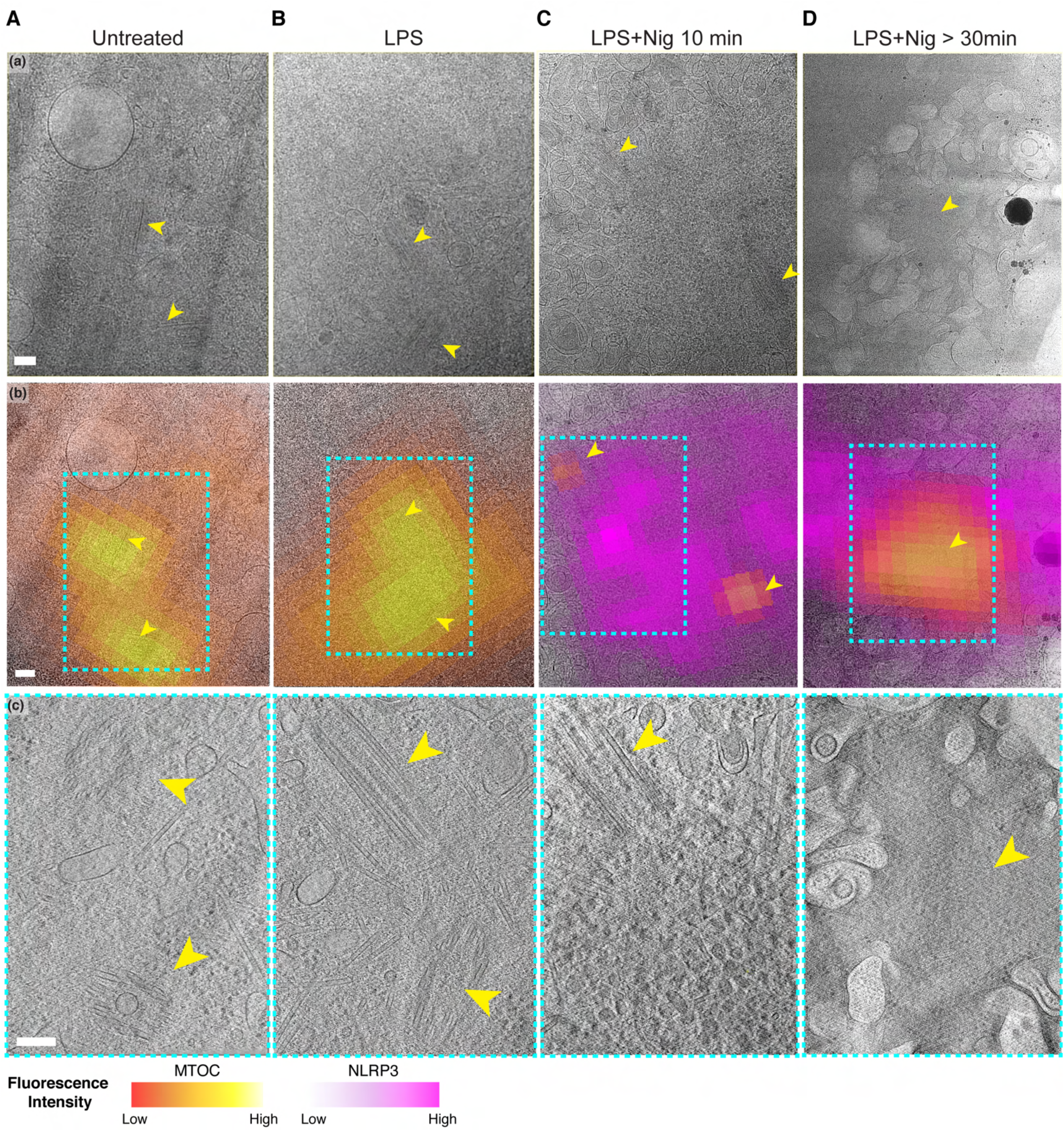
Progression of NLRP3 inflammasome activation. **(A-D)** Representative tomographic slices with increasing magnification showing the MTOC and formation of the NLRP3 inflammasome condensate ordered by chronological sequence with **(a)** cryoET atlas, **(b)** fluorescence correlated cryoET atlas overlaid with (a), **(c)** representative tomographic slices obtained from designated imaging area indicated as cyan dashed lines in (b). The MTOC and NLRP3 inflammasome are colored in yellow and magenta respectively based on fluorescence intensity profile. Two centrioles from the MTOC are indicated as yellow arrows. Number of incidents at each stage: No activation, N=12; LPS, N=10; LPS+Nig 10-20 min, N=10; LPS+Nig >30min, N=3. (scale bar = 200nm for all panels).

**Fig. 2.**
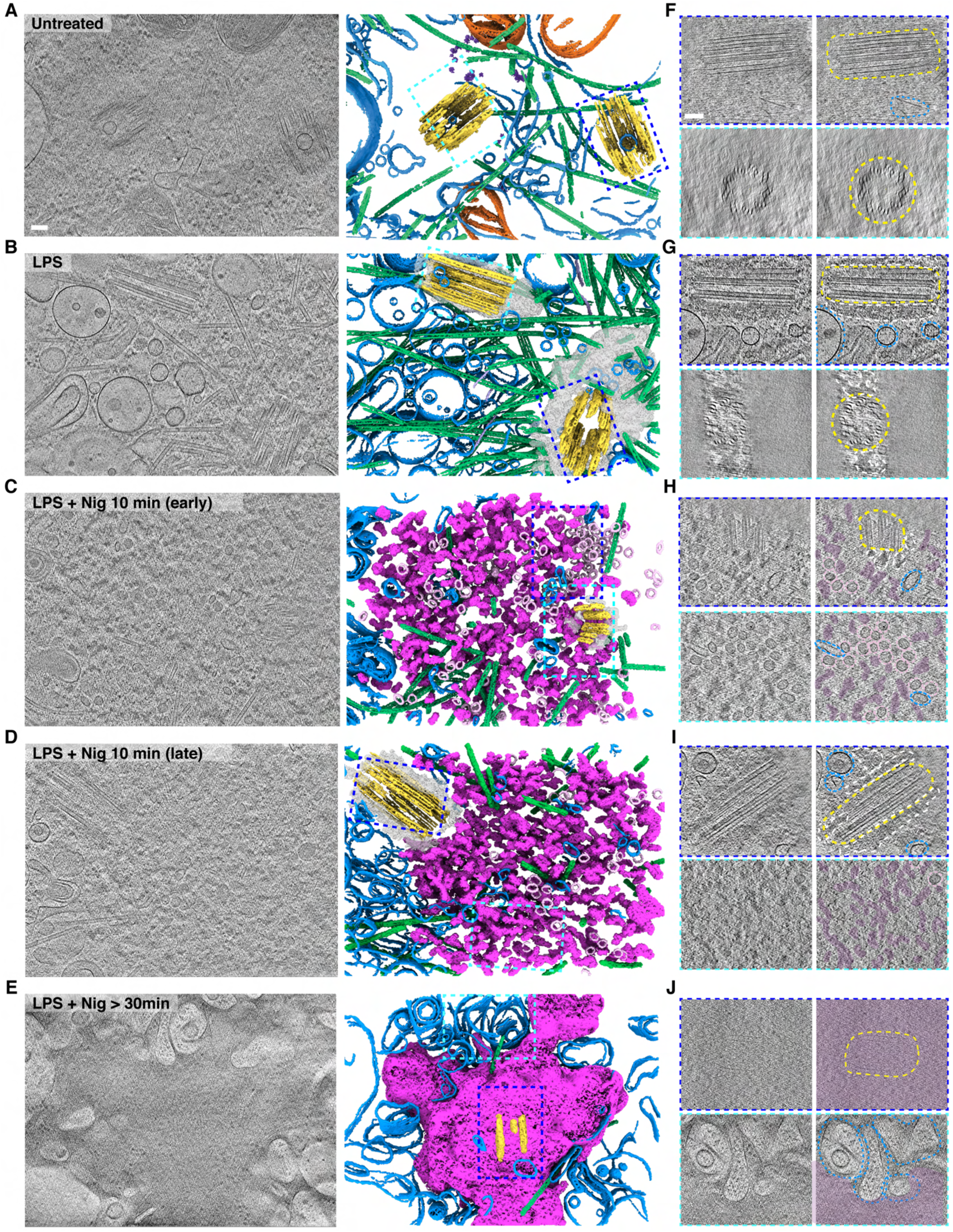
NLRP3 inflammasome activation leads to formation of condensate at the MTOC. **(A-E)** Representative segmented models and tomographic slices illustrating NLRP3 inflammasome activation (MTOC, yellow; microtubules, green; NLRP3 associated vesicles, light pink; NLRP3 condensate, magenta; membrane, blue; mitochondria, orange, scale bar = 100 nm). **(F-J)** Close-up view highlighting cellular features observed in tomographic slices in A-E with an illustrative view. Origins of insets are indicated in segmented models.

**Fig. 3.**
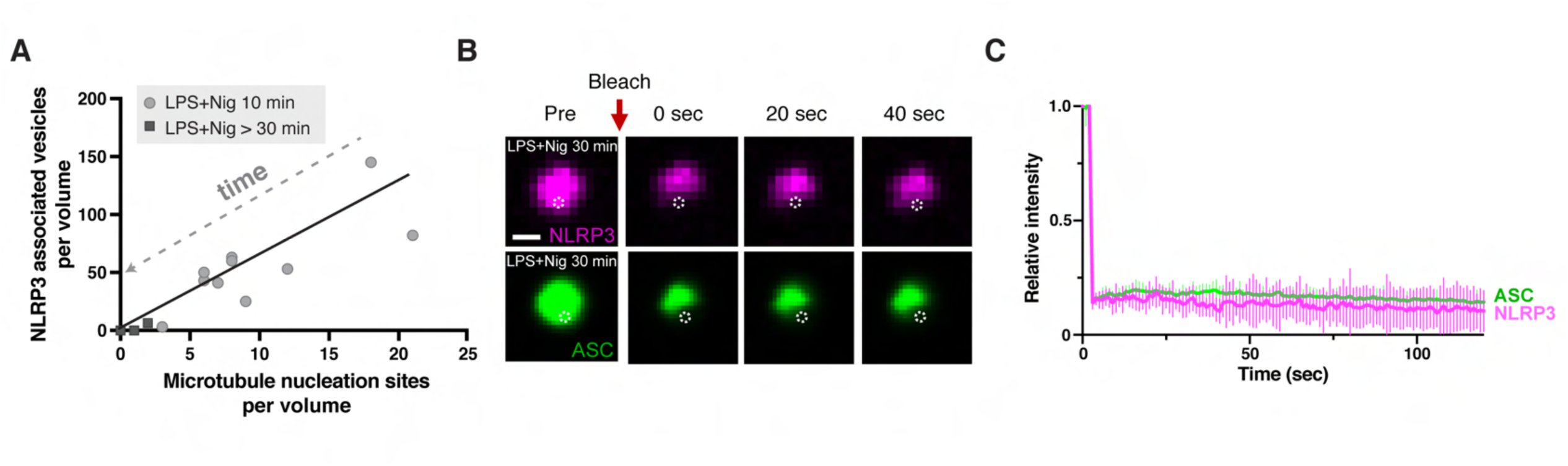
NLRP3 inflammasome becomes an immobilized solid condensate. **(A)** Correlation analysis of number of NLRP3 associated vesicles and number of microtubule nucleation sites within NLRP3 condensate. Each data point represents one cell. Cells treated with LPS+Nig 10 min are indicated as grey circles; cells treated with LPS+Nig > 30 min are marked as dark grey squares**. (B-C)** Tracking of NLRP3 mobility by fluorescence recovery after photobleaching (FRAP). Bleached areas indicated by white dashed circle (scale bar=5µm).

In order to examine the dynamics of NLRP3 within the condensate, we performed fluorescence recovery after photobleaching (FRAP) experiments using mSL-NLRP3 cells also expressing mNeonGreen-ASC. Neither NLRP3 nor ASC in the inflammasome speck recovered after photobleaching (**Fig. 3B-C**), suggesting the condensate had solidified.

### NLRP3 inflammasome priming and activation perturbs the MTOC

In our tomographic reconstructions, each centriole was seen to be a ring of nine triplets of microtubules (MTs) 500-700 nm in length and approximately 250 nm in diameter (**Fig. 2F-G, I**), consistent with previous studies (35–38). In untreated cells, the two centrioles were found to lie perpendicular to each other, spaced on average 500 nm apart (**Fig. 4A**), also consistent with other studies (39). As these distances are close to the diffraction limit of our imaging system, under these conditions the two centrioles’ fluorescence signal partially overlapped (**Fig. 1A-B**).

**Fig. 4.**
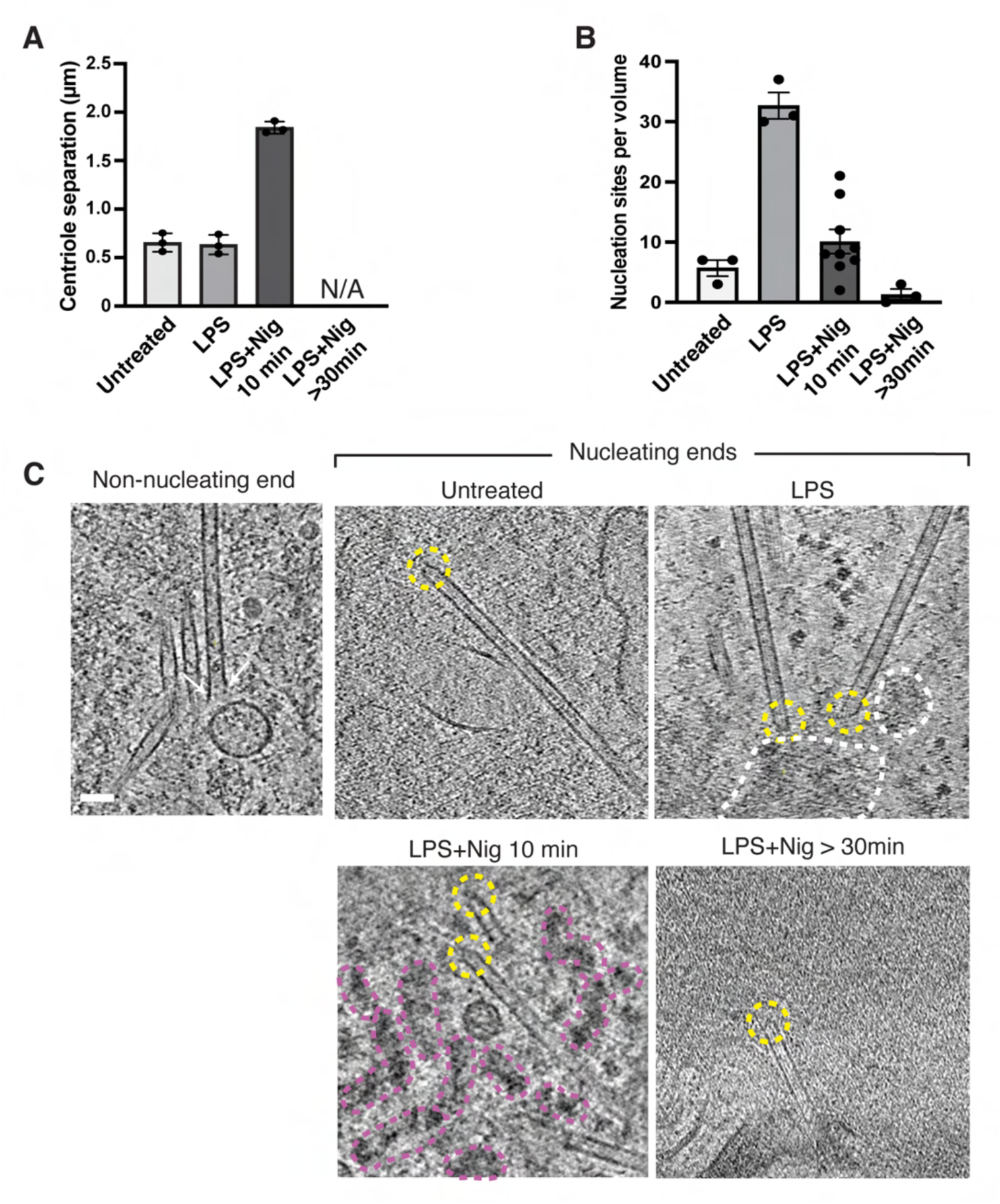
NLRP3 inflammasome activation alters MTOC conformation. **(A)** Distance of centriole separation in response to NLRP3 inflammasome activation (one-way ANOVA, p<0.0001). **(B)** Quantification of the number of nucleation sites in presence of the complete MTOC. Each data point represents one cell (one-way ANOVA, p<0.0001). (**C**) Representative tomographic slices showing non-nucleating ends and MT nucleation sites at different conditions. The non-nucleating end is marked as two white arrows and the nucleation site is circled in yellow; PCM and NLRP3 condensate are indicated as white and magenta, respectively (scale bar=50nm).

The pericentriolar material (PCM) has been described as a phase-separated compartment in close contact with the centrosome that acts as a scaffold for nucleating microtubules (40, 41). While we could not discern PCM in the cryo-tomograms of untreated cells, priming with LPS resulted in the appearance of dark amorphous densities with well-defined boundaries, tightly associated with the centrioles (**Fig. 2B, G, Fig. S3A**). As a similarly-dark amorphous material was identified as PCM in a previous report (38), we interpret this material to be PCM. In untreated cells, we observed only a few microtubule nucleation sites in close proximity to the centrioles, but in LPS-primed cells there were many more, located within the PCM (**Fig. 4B-C**). Microtubule nucleation sites were identified by conically-shaped γ-tubulin ring complexes (γtuRC) at the tip (**Fig. 4C**) (42–44). Recruitment of PCM and enhanced microtubule nucleation are both recognized hallmarks of interphase centrosome maturation (45). Thus, our observations infer that NLRP3 inflammasome priming induces features of centrosome maturation.

Ten minutes after nigericin stimulation, the two centrioles were observed further apart, approximately 2 µm away from each other (**Fig. 4A**), and nearly parallel (**Fig. 1C**). Notably, before milling, the SiR (tubulin) fluorescence appeared as a single punctum, but after milling, each centriole could be resolved (**Fig. S2**). The distance between centrioles has been reported to be cell-cycle dependent. Fully separated centrioles (>2 µm) are formed at late G2 phase (46–48). Thus, NLRP3 inflammasome activation induces changes characteristic of cell cycle progression.

### Inflammasome activation induces Golgi cisternae swelling and disruption

In untreated cells, we observed Golgi stacks with 20-30 nm thick cisternae, similar to what has been reported in previous studies (49) (**Fig. 5A-B**). Following LPS-priming, the diameter of cisternae expanded to 80-100 nm (**Fig. 5A-B**), an effect previously inferred to reflect increased cargo loading (50, 51). To investigate Golgi structure with a wider field of view than possible with cryo-ET, we stably expressed TGN38-mNeonGreen and recorded fluorescence images. Indeed, area positive for TGN38 also appeared to be larger in LPS-primed cells compared to untreated cells, suggesting Golgi expansion (**Fig. 5C**). We hypothesize therefore that the reported increase in NLRP3 recruitment to TGN by LPS priming (52).

**Fig. 5.**
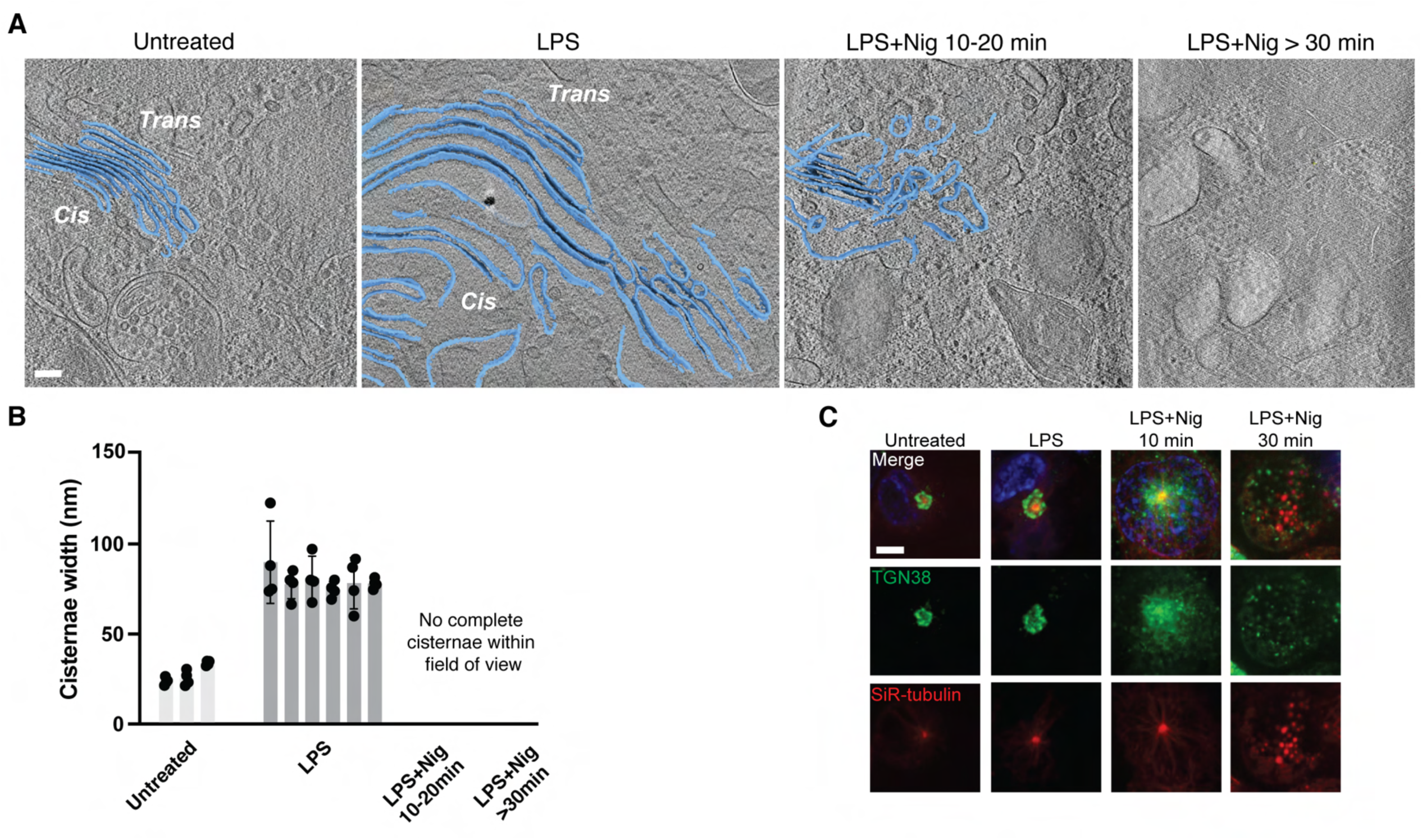
Nigericin stimulation disrupts Golgi integrity. **(A)** Overlaid segmented Golgi structures on tomographic slices (Golgi cisternae, blue; z=12nm, scale bar=100 nm). **(B)** Quantification of the cisternae spacing in response to LPS. Each bar graph represents 3 distance measurements for each Golgi structure. Face of the Golgi is assigned based on relative distance to the nucleus on lamellae where the *Cis* face is closer to the nucleus. All cells come from different passages and different grids (no activation, n=3; LPS, n=6; one-way ANOVA, p<0.0001). **(C)** Live cell imaging of mScarlet-NLRP3, TGN38-mNeonGreen and SiR-tubulin before and after NLRP3 inflammasome activation (scale bar=5 µm).

In LPS-primed cells we also noticed clusters of ∼50-nm vesicles close to the swollen Golgi (**Fig. 6A, Fig. S4A-C**). While the vesicles appeared to contain some protein density, no regular arrangement could be discerned (**Fig. S4B**). The clusters’ positions and vesicle concentration correlated with mScarlet fluorescence intensity, indicating the presence of concentrated NLRP3 (**Fig. S5**). Similarly-sized vesicles were not observed in untreated cells or regions outside NLRP3 fluorescence (**Fig. 6A**). The presence of these vesicles in LPS-primed cells is consistent with the recent finding that NLRP3 is immobilized on Golgi before addition of nigericin (51). In nigericin-stimulated cells we again saw 50-nm vesicles, in larger numbers, but we could not detect any special spatial correlation between the clusters of vesicles and mScarlet fluorescence because the fluorescence signal was uniformly high across the entire area due to the growth of the condensate (**Fig. S5**).

**Fig. 6.**
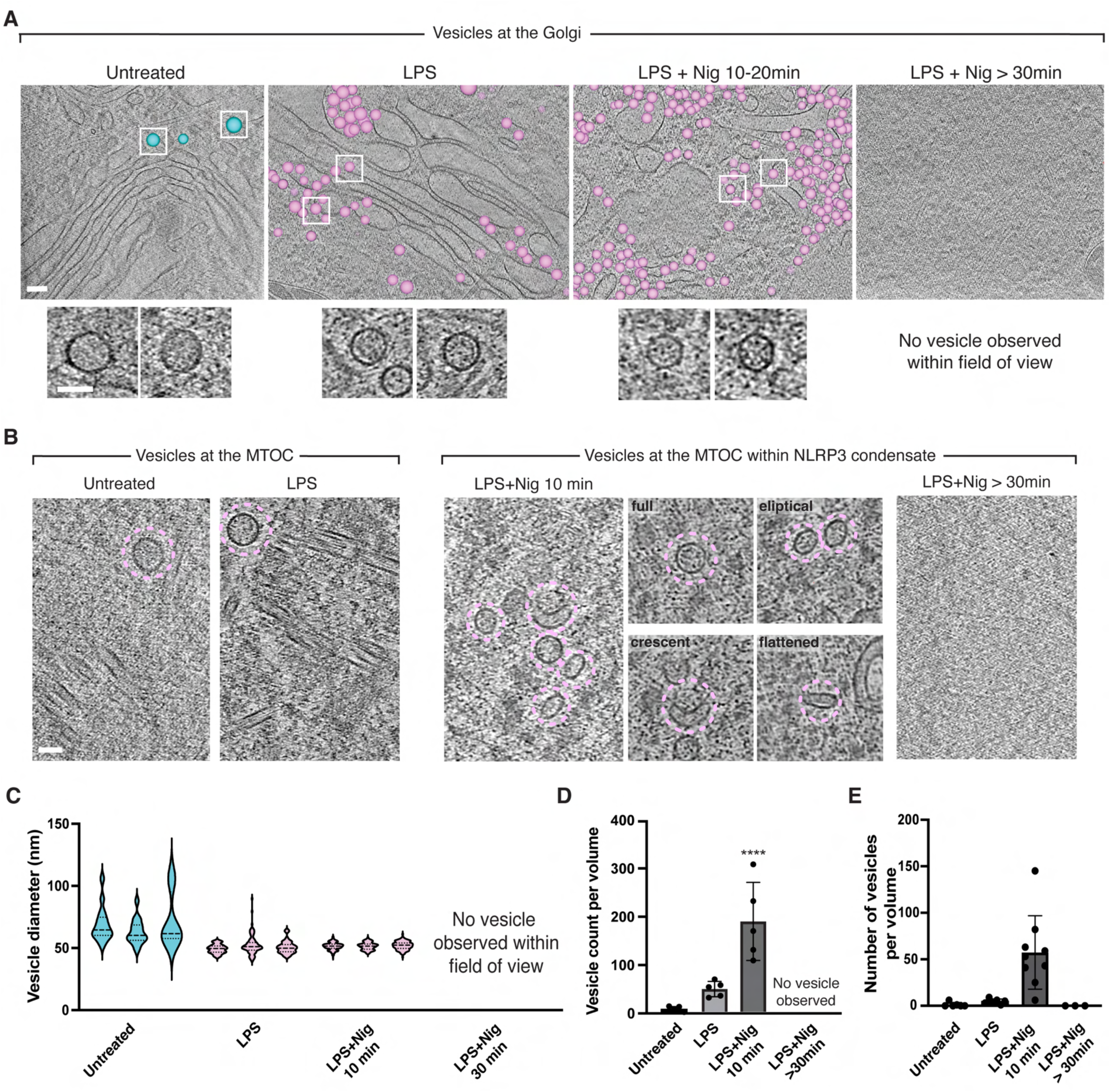
NLRP3-associated vesicles trafficking to the MTOC independent of retrograde transport. **(A)** Overlaid segmented model on tomographic slices showing naturally secreting vesicles in the cytosol (untreated) and NLRP3 associated vesicles (LPS, LPS+Nig 10 min, LPS+Nig > 30 min) (naturally secreting vesicles, cyan; NLRP3-associated veislces, light pink; scale bar=100 nm). Close-up view of representative vesicles observed under each condition was indicated as white boxes and showed in insets (scale bar=50 nm). **(B)** Representative tomographic slices showing vesicles near the MTOC before and after NLRP3 inflammasome condensate formation (z=12nm). Vesicles are circled in light pink. **(C)** Quantification of vesicle diameters in response to NLRP3 inflammasome activation. Each violin plot represents one cell (one-way ANOVA, p<0.0001). **(D)** Quantification of vesicle numbers observed near the Golgi in response to NLRP3 inflammasome (one-way ANOVA, p<0.0001). Each data point represents one cell. **(E)** Quantification of number of vesicles present at the MTOC at different conditions. Each data point represents one cell (one-way ANOVA, p<0.0001).

After nigericin stimulation, little or no intact Golgi structure was seen (**Fig. 5A**). Instead, we observed diverse irregular-shaped membrane structures (**Fig. 5A**). To investigate Golgi structure with a wider field of view than possible with cryo-ET, Using THP-1 cells stably expressing TGN38-mNeonGreen, the TGN38 fluorescence appeared as dispersed speckles after nigericin stimulation (**Fig. 5C**), consistent with the cryo-ET data. We hypothesize that the 50-nm vesicles are derived from the Golgi during its dispersion (19, 53) and carry NLRP3 to the MTOC (**Fig. 6A-B**). Consistent with this hypothesis, some vesicles were non-spherical or even flattened, as expected for different stages of content release (**Fig. 6B**).

To our surprise we found very few 50-nm vesicles in close contact with microtubule networks or intermediate filaments (**Fig. S4D**). In mouse macrophages, NLRP3 has been demonstrated to interact with microtubules and intermediate filaments and inflammasome activation depends on HDAC6-mediated microtubule transport (14, 16, 54, 55). We therefore tested whether inflammasome activation was also dependent on HDAC6-mediated microtubule transport in our human cells. We found that it was not in both WT and mSL-NLRP3 cells, as shown by insensitivity to the microtubule disrupter colchicine and the HDAC6 inhibitor tubacin (**Fig. S1C-D**). This microtubule-independence is in keeping with the recently shown NEK7 independence of NLRP3 activation in human macrophages (52).

Notably, we imaged a few LPS-primed cells in which both the MTOC and Golgi stacks were retained in the final lamellae. In these cases the swollen Golgi stacks appeared as close as 200 nm away from a centriole (**Fig. S4A**). By contrast, in untreated mSL-NLRP3 THP-1 cells the MTOC and Golgi were never seen together in the same cryo-tomogram, indicating they are more than a few microns apart (the field of view of one tomogram), and suggesting that inflammasome activation might bring the entire Golgi close to the MTOC, obviating the need for microtubule-based directed transport of vesicles across long distances.

### Mitochondria are recruited to inflammasomes and appear damaged

Mitochondria have been reported to play a role in the NLRP3 inflammasome activation pathway (56–59). A recent study showed abnormal cristae structure in nigericin-stimulated ASC-overexpressing mouse iBMDMs (23). We observed a plethora of mitochondrial morphologies in untreated, LPS-primed, and early nigericin-stimulated (10 min) cells, predominantly displaying intact outer membranes and evenly distributed cristae, with rare cases showing loss of cristae or irregular-shaped cristae (**Fig. S6A**).

However, the number of mitochondria within 5 µm of the MTOC and NLRP3 condensate increased dramatically after 10 min of nigericin treatment (**Fig. 7A-B**). Cells treated with nigericin for more than 30 min revealed much larger mitochondria with reduced cristae (**Fig. S6A**), an indication of damage (60). Mitochondria were seen in clusters, and some appeared to be fusing (**Fig. 7B, Fig. S6B**). The formation of mitochondrial clusters was previously reported in cells overexpressing mitochondrial fusion protein Mfn2, which ultimately underwent caspase-mediated apoptosis (61, 62). Increased mitochondrial clustering and fusion have also been observed in cells with elevated levels of reactive oxygen species (ROS) (63, 64). As GSDMD-mediated mitochondrial damage has been shown to induce ROS (65, 66), we speculate that the mitochondrial clustering and fusion we observed here are caused by similar pathways. After 10 minutes of nigericin stimulation, mitochondria were also seen in direct contact with autophagosomal-like structures surrounding the NLRP3 condensate (**Fig. 7C; Fig. S7A-B**). We previously showed that NLRP3 inflammasome puncta colocalize with the autophagy marker LC3b (14).

**Fig. 7.**
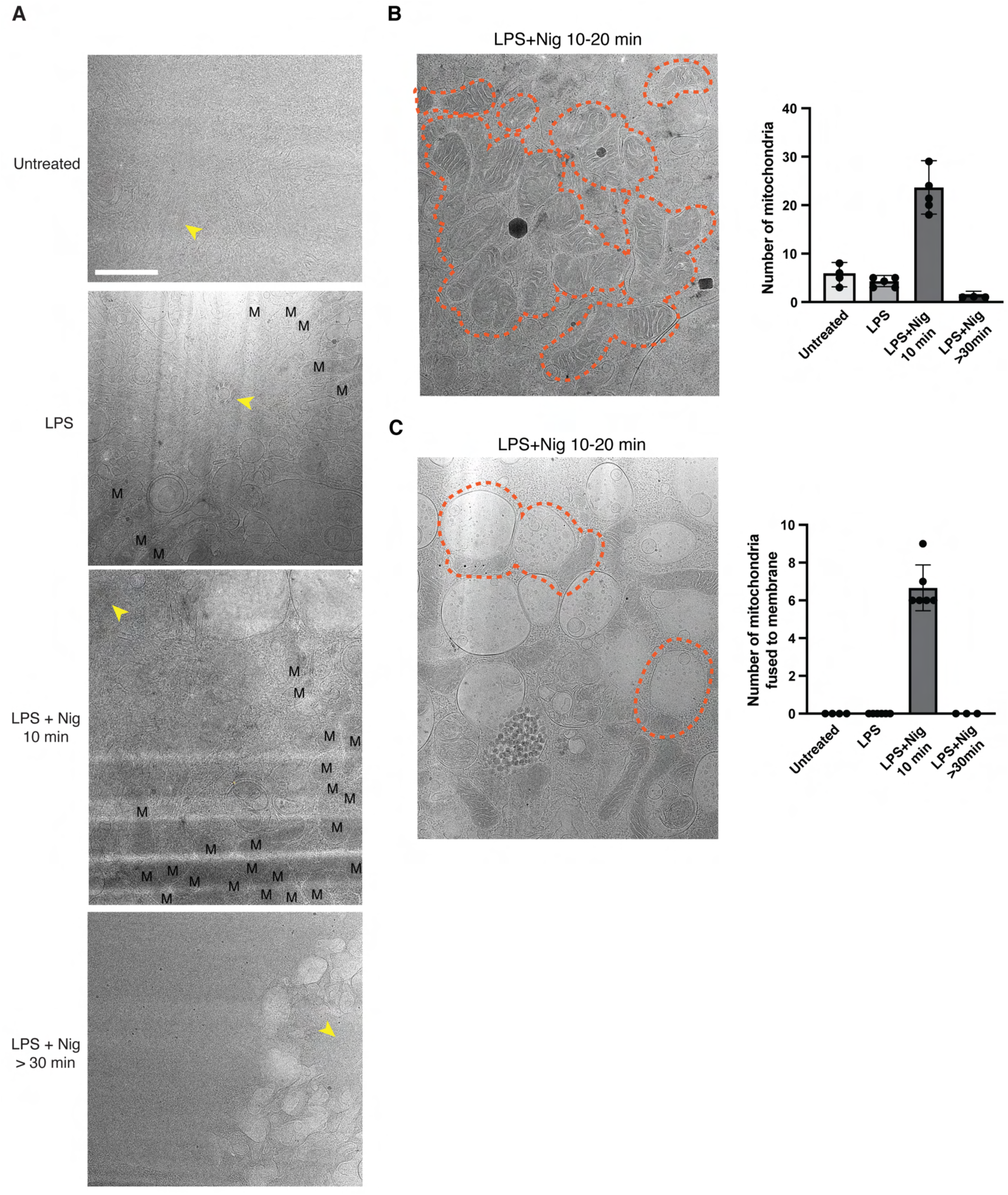
NLRP3 inflammasome activation induces changes of mitochondria. **(A)** Representative atlases of the MTOC and mitochondria in proximity (scale bar=1 µm). Centrioles are marked in yellow and mitochondria are denoted as “M”. **(B)** Representative clustered area is denoted as dashed orange area. Quantification of number of mitochondria present within 5 µm from the MTOC. Each data point represents one cell (one-way ANOVA, p<0.0001). **(C)** Quantification of the amount of mitochondria with close contact to autophagic membranes (one-way ANOVA, p<0.0001). Representative examples showing mitochondria with close contact to autophagic membranes are indicated as dashed orange area.

Here we confirmed that NLRP3 fluorescence again colocalized with LC3b in these cells, indicating that at least some of the structures were autophagic (**Fig. S7C**).

## Discussion

### Mechanism of NLRP3 inflammasome activation

Our data support the following model for NLRP3 inflammasome activation (**Fig. 8**). LPS priming enhances the Golgi recruitment of NLRP3, causing Golgi cisternae swelling (50, 51). Swollen Golgi cisternae give rise to special 50-nm vesicles loaded with NLRP3 and perhaps other inflammasome material. At the MTOC, LPS priming leads to increased PCM density and microtubule nucleation (**Fig. 8**). The Golgi and the MTOC move closer together. Nigericin stimulation then leads to complete Golgi dispersion and the 50-nm vesicles diffuse the short distance to the MTOC without active transport and unload their contents. These contents aggregate and recruit/concentrate ASC and caspase-1, leading to caspase-1 activation. Mitochondria are recruited and damaged, and autophagasomes appear. The condensate eventually solidifies (no exchange of materials), and pyroptosis is triggered.

**Fig. 8.**
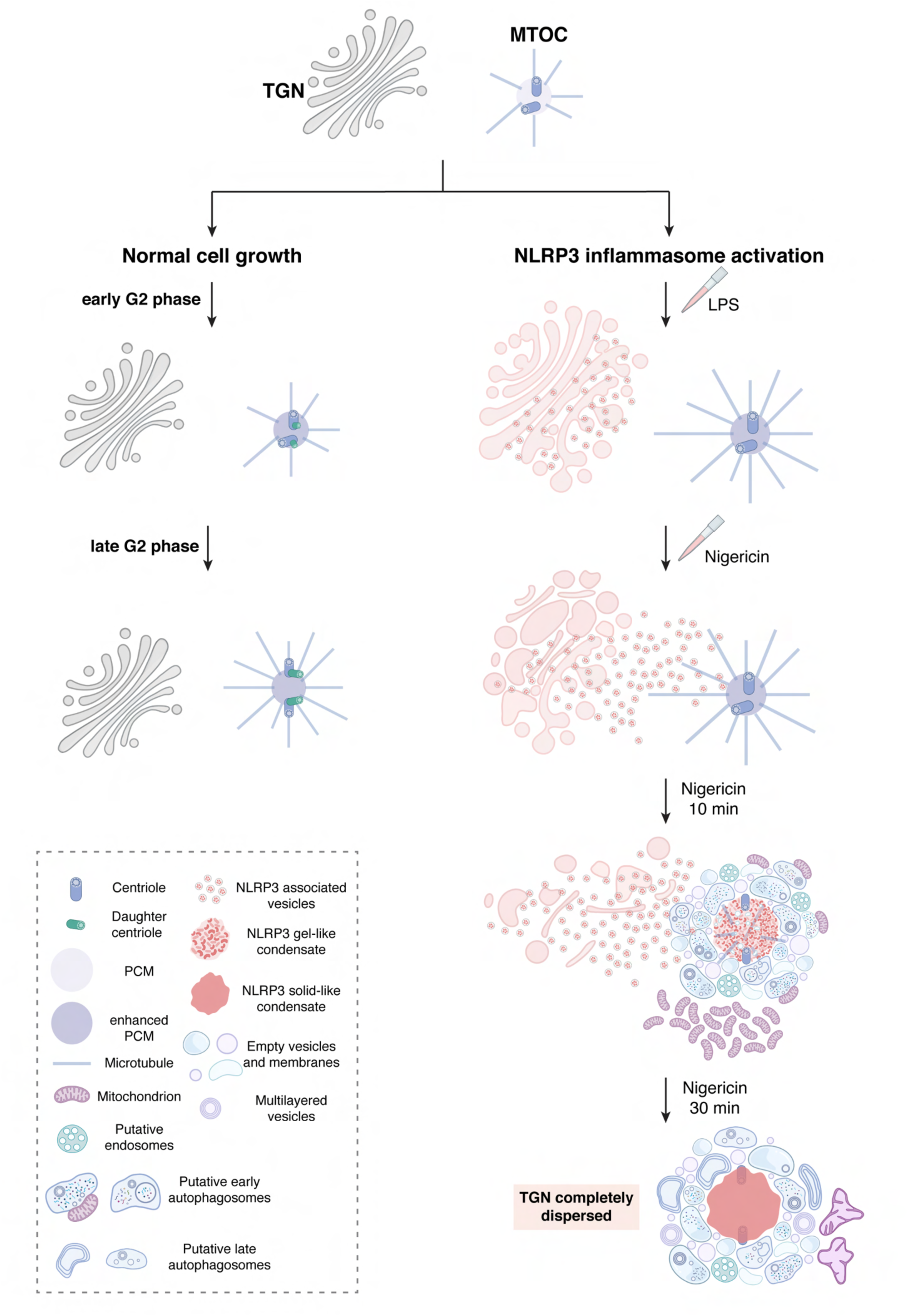
Progression of NLRP3 inflammasome activation at the MTOC. Cartoon representations are annotated in the bottom left. The NLRP3-associated vesicles were modeled based on previous structures (PDB: 7FLH) (19). [Figure was created with BioRender (https://BioRender.com).]

We were unable to identify regular, ordered structures within the NLRP3-containing patches or later condensates. This observation rules out the presence of long, extended polymer tubes, but not smaller complexes such as the partial inflammasome disks seen in recombinant samples (10), which might have been present. Importantly, regular, ordered structures were also not discernable in single-particle images of those recombinant samples - the disc shape and structure only became apparent after extensive 2D classification, which required hundreds of thousands of particles (orders of magnitude more than we have in the cryo-tomograms). When in a related study ASC was over-expressed, obvious filament structures were observed in situ (23). Since NLRP3 inflammasome activation does not require ASC overexpression, we posit that extended filamentous structures, which were not observed in our tomographic reconstructions, are not necessary for NLRP3 inflammasome activation. Instead, smaller assemblies such as partial discs must be sufficient to drive dimerization of the caspase domain of caspase-1 and proximity-induced enzymatic activation (10).

### Inflammation and cell division

Our results suggest why formation of the NLRP3 inflammasome and normal cell division are mutually exclusive (67): we found that LPS priming led to increased PCM and microtubule nucleation, but NLRP3 aggregate formation disrupted centriole spacing and created a condensate that enveloped the MTOC likely limiting its accessibility by other cell division cofactors (45). These observations are consistent with two recent findings that microtubule nucleation requires dynamic PCM (41), and loss of PCM has been reported as a consequence of NLRP3 inflammasome activation (68).

### Phase separation as a general mechanism to promote innate immune signaling

Biomolecular condensate formation by phase separation has been implicated in numerous biological processes (69–71) including immunity to concentrate proteins and drive rapid, switch-like signaling (72–76). Some of us (Shen et al.) previously found that NLRP6, an inflammasome protein in the same family as NLRP3, contains intrinsically disordered regions and forms dynamic condensates when bound to viral dsRNA or bacterial lipoteichoic acid (74). The condensate can recruit the ASC adaptor, which in turn recruits caspase-1, leading to solidification of the condensate (74). Our results suggest a similar mechanism for the NLRP3 inflammasome. The phase separation and multi-protein complexes likely both facilitate threshold behavior, spatial control and raised local concentration (77–80). Further elucidation of these mechanisms could open up new avenues for therapeutic strategies to dampen overactive immune pathways.

### Analogy to short-distance presynaptic vesicle trafficking

Intracellular vesicle trafficking is a well-known process for exchanging materials between membrane-enclosed organelles (81). Facilitated by motor proteins and cytoskeleton tracks, cargoes secured in vesicles can be transported long distances from one cellular compartment to another. However, less is known about vesicle trafficking across short distances. One example occurs at the synapse where presynaptic vesicles in the phase-separated reserve pool travel approximately 200-300 nm to join the phase-separated active zone (82–85). Active transport is apparently not required. The NLRP3-associated vesicles observed in our study share similar features as presynaptic vesicles: both are spherical vesicles of on average 50 nm in size, enriched in high abundance at particular subcellular regions, and diffuse short distances to join a phase-separated compartment (86, 87).

### Flourescence-guided cryo-FIB milling

Our tri-coincident cryo-FIB-SEM-flourescence instrument allowed us to routinely uncover and preserve in lamella sub-diffraction-limited fluorescent targets like centrioles and nascent inflammasomes. The resolution of the fluorescence microscopy on the final polished lamella was even sufficient to correlate with specific clusters of vesicles and their concentration, providing strong evidence that the vesicles contained NLRP3. By freezing cells at different time points, we were able to elucidate the entire inflammasome activation process. When comparing different kinds of experiments, however, timing may be different in cells grown on EM grids, glass slides, petri dishes, or other contexts. Collectively, these instrumental and methodological advances open new opportunities for directly visualizing molecular complexes *in situ* by cryo-ET.

## Acknowledgments

We express our gratitude to Delmic B.V. for development of the tri-coincident cryo-FIB-SEM platform. We would like to acknowledge the Stanford-SLAC Cryo-EM Center and Stanford University Cryo-electron Microscopy Center (cEMc) for instrumentation with special thanks to Chensong Zhang and Lydia-Marie Joubert. We thank the Core for Imaging Technology & Education (CITE) and Microscopy Resources On the North Quad (MicRoN) at Harvard Medical School for help with light microscopy. We also thank Caltech Cryo-EM Facility for instrumentation at the initial phase of this project. This work was supported in part by National Institutes of Health (AI127401 to GJJ), Grant 2021-234593 from the Chan Zukerberg Initiative DAF (to PDD), advised fund of Silicon Valley Community Foundation (to PDD), in part by the Panofsky Fellowship at the SLAC National Accelerator Laboratory as part of the Department of Energy Laboratory Directed Research and Development program under contract DE-AC02-76SF00515 (to PDD), National Institutes of Health (AI77778 to HW), and Cancer Research Institute Postdoctoral Fellowship (to MW).

## Materials and Methods

### Constructs and cloning

pLenti-CMV-Flag-mScarlet-NLRP3 was previously described (19). TGN38-mNeonGreen and mNeonGreen-ASC were cloned into the pHAGE-EF1⍺ vector between XbaI and NheI sites using Gibson Assembly Master Mix (NEB, Cat. No: M5510).

### Generation of stable cell lines

To produce lentiviral particles, HEK293T cells (80% confluence) in 10 cm dishes were co-transfected with 10 μg pLenti-CMV-Flag-mScarlet-NLRP3, 10 μg pHAGE-EF1⍺-TGN38-mNeonGreen, or 10 μg pHAGE-EF1⍺-mNeonGreen-ASC, and packaging plamids 7.5 μg of psPAX2 and 3 μg pMD2.G (Addgene plasmids #12260 and #12259). 12 hours after transfection, the media were removed and replenished with 8 mL of fresh medium. The supernatants containing lentiviral particles were harvested twice at 48 and 72 hours after transfection, filtered through 0.45 μm filter (Pall Corporation, Cat. No: 4184), concentrated using Lenti-X™ Concentrator (Takara Bio, Cat. No: 631231) and stored at -80 °C until use.

To infect THP-1 cells with lentiviruses, cells were cultured in media containing lentiviruses and 20 μg/mL DEAE-Dextran (Sigma-Aldrich, Cat. No: 93556). To increase the transduction efficiency, fluorescence-activated cell sorting (FACS) was performed using a FACSAria II cell sorter from Becton Dickinson equipped with FacsDiva version 8.03. The instrument was set up with a 100 μm nozzle. The sorted populations were gated to exclude double, dead and autofluorescent cells. The cells were recovered for at least 7 days before performing subsequent experiments.

### Cell culture

Human monocytic THP-1 cells were maintained in Roswell Park Memorial Institute (RPMI) 1640 Medium (Thermo Fisher Scientific, Cat. No: 11875085), supplemented with heat inactivated 10% FBS and 0.05 mM 2-mercaptoethanol (Thermo Fisher Scientific, Cat. No: 31350010),10 mM HEPES (Thermo Fisher Scientific, Cat. No: 15630080), 1 mM Sodium Pyruvate (Thermo Fisher Scientific, Cat. No: 11360070), 100 units/mL of penicillin and 100 μg/mL of streptomycin.

The NLRP3 KO THP-1 cells were purchased from company (Invivogen, Cat. No: thp-konlrp3z). NLRP3 KO THP-1 reconstituted with mScarlet-NLRP3 cells were generated via lentivirus infection and flow cytometry sorting as described above, which is used for Cryo-electron tomography throughout the manuscript.

All cells were maintained at 37 °C with 5% CO_2_. For inflammasome activation studies, THP-1 cells were differentiated for 48 hours with 100 nM phorbol myristate acetate (PMA, Sigma-Aldrich, Cat. No: P8139– 5MG), followed by priming for 4 hours with 1 μg/mL lipopolysaccharides (LPS, Invivogen, Cat. No: tlrl-b5lps) before stimulation with 20 μM nigericin (Sigma-Aldrich, Cat. No: N7143-10MG) for the indicate times. When needed, 10 μM microtubule polymerization inhibitor Colchicine (Sigma-Aldrich, Cat. No: C9754–100MG), or 20 μM HDAC6 inhibitor Tubacin (Sigma-Aldrich, Cat. No: SML0065-1MG) were used for 2 hours before nigericin treatment.

### Sample preparation for cryoET

mScarlet-THP1 cells were maintained as described in previous section. Cells were differentiated for 24 hours in tissue culture treated plate and seeded onto human fibronectin (25μg/mL in 1xPBS, Advanced Biomatrix, Cat. No: 5050) coated grids (R2/2, Au 200-mesh London Finder grid coated with extra thick carbon, Electron Microscopy Sciences) for another 24 hours to allow sufficient adherence. On the day of plunge freezing, cells on grids were primed with LPS (1 μg/mL) and stained by SiR-tubulin (1 μM, Cytoskeleton, Inc. Cat. No: CY-SC002) for 4 hours (88). Cells in the treatment group were further treated with 20 μM nigericin for NLRP3 inflammasome activation for 10-20min or 30-45min. Untreated cells were stained by SiR-tubulin only for 4 hours. Grids were plunge-frozen in liquid ethane (Airgas) at different time points on a MarkIV vitrobot (Thermo Fisher Scientific) at 100% humidity with manual blotting. Each condition was frozen based on the following order: after 4 hours for both untreated and LPS treated cells; 4 hours plus 10-20min for LPS+Nig 10-20 min treated cells; 4 hours plus 30-45min for LPS+Nig >30 min treated cells.

### In-situ fluorescence guided cryoFIB-SEM milling

Frozen grids were clipped into AutoGrids with 4 milling slots (Thermo Fisher Scientific) and loaded onto the Ariyscan laser scanning microscope (Zeiss LSM880) equipped with a cryogenic sample stage (Linkam, CMS196) for inspection of colocalized NLRP3 and MTOC puncta and ice quality. NLRP3 signal was detected by excitation at 555 nm. MTOC signal was detected by excitation at 625 nm. Grids were then transferred to an Aquilos2 Dual-Beam system (Thermo Fisher Scientific) with customized integrated fluorescent light microscope (ENZEL from Delmic) that is coincident with the FIB and SEM of the Aquilos2 as described previously (29). Organo-platinum (Pt) deposition was applied to grids via the gas injection system (GIS) for 20-30 seconds. The NLRP3 inflammasome was identified as the colocalized NLRP3 and MTOC puncta observed on the ENZEL system equipped with a 100x long working distance vacuum objective (NA 0.85). Materials above and below the target of interests were ablated accordingly monitored by real-time fluorescence. The stepwise milling was performed as described in previous studies (89, 90). In brief, rough milling, thinning, and polishing were done at 0.1nA, 30pA, 10pA at 30kV. Final fluorescent images were taken when the thickness of the lamella was approximately 200nm with the following imaging parameters: NLRP3, ∼0.96 W/cm^2^ of 550 ± 10 nm with 1.5-2 seconds exposure; MTOC, ∼ 0.46 W/cm^2^ of 625 ± 10 nm with 1-2 seconds exposure; Reflected brightfield, ∼0.1 W/cm^2^ of 550 ± 10 nm with 500 milliseconds exposure.

### Tilt series acquisition guided by correlated fluorescence microscopy

Atlases and tilt series were acquired on a Titan Krios G2 (Thermo Fisher Scientific) at 300 keV equipped with a K3 Summit detector and BioQuantum post-column energy filter (Gatan) operated in counting and dose fractionation mode with 20 eV slit in. Atlases were collected as joint serial sections at 6,500x magnification under low dose mode in SerialEM software suite (91, 92). Tilt series were collected at a pixel size of 3.465 Å with a total dose of approximately 120-160 e^-^/Å^2^, ranging from -48° to 60° to account for a ∼6° milling angle with 2-degree increment using SerialEM. This study included a total of 52 tilt series: 11, 17, 20, 4 for untreated, LPS, LPS+Nig 10-20min, LPS+Nig >30min, respectively.

### Tomogram reconstruction and quantification of cellular features

Unbinned image frames were preprocessed in WARP with patch motion correction, CTF estimation, and defocus estimation, which ultimately assembled to aligned tilt series with a binning factor of 2 (93, 94). Aligned tilt series were reconstructed manually in IMOD at a pixel size of 13.86 Å with 30 iterations of SIRT-like filter following CTF correction (95, 96). Tomogram reconstructions containing TGN were loaded in FIJI for measuring cisternae width. Tomogram reconstructions containing NLRP3 associated vesicles were loaded in IMOD (95, 96). The center of each vesicle was manually set and used the size function to measure the radius of each vesicle in pixel, which later was converted to vesicle diameter in nanometers.

### Tomogram segmentation and visualization

Tomograms used for segmentation were imported to Dragonfly 2022.2 and subject to Gaussian filtering to enhance image contrast. Training data sets were prepared by manually segmented several slices containing microtubules, large membrane structure, Golgi stacks, small vesicles, ribosome, and mitochondria. The Neural Network Model was generated with a dimension of 2.5D and 3 slices. NLRP3 condensate and PCM were segmented using the thresholding tool. Segmented features were exported independently and later processed in FIJI with Gaussian blurring (sigma=0.8) and imported to ChimeraX (version 1.7.1) for visualization (97). Tomograms containing NLRP3 vesicles were manually segmented in IMOD for microtubules, NLRP3 associated vesicles, and intermediate filaments.

### Correlated fluorescence microscopy

Registration of fluorescence and electron tomography data was performed utilizing a customized toolkit compatible with MATLAB version R2023b. Briefly, multi-channel fluorescent images obtained from the ENZEL system were firstly correlated to atlases collected at 6500x magnification using obvious features (e.g. lamellae edges, visible ice contamination, or grid holes) present in both brightfield optical microscopy and the electron microscopy. A total of 8-10 pairs of reference points were selected accordingly for precise correlation. These point pairs were used to calculate a projective transformation to carry the fluorescence data to this low magnification EM space. Secondly, a new set of points visible in the registered atlases and in a z-projection of the tomographic reconstruction were selected. These points largely included visible cellular features (e.g. large membrane structure, vesicles, mitochondria, or MTOC). A total of 10-13 pairs of reference points were manually selected for registration and the computation of a similarity transformation to carry the atlas space into the tomography space. Lastly, the projective and similar transformation were serially applied to the fluorescence images to bring the fluorescence images to the tomography space. For visualization, final correlated images containing SiR-tubulin and NLRP3 channels were generated by overlaying fluorescent channels with actual tomogram reconstruction (z=12nm).

### Vesicle correlation analysis

Correlated atlases and tomogram reconstructions were computed as described in previous section. Vesicle x and y coordinates were extracted from IMOD segmented models and later correlated with registered tomogram reconstructions. Number of vesicles within each registered fluorescent pixel was plotted and fitted to a simple linear regression.

### LDH cytotoxicity assay

WT THP-1, NLRP3 KO THP-1 and NLRP3 KO THP-1 reconstituted with mScarlet-NLRP3 were seeded on a 24 well plate. After differentiation, priming and NLRP3 activation, cell supernatants were analyzed for LDH activity using LDH-Glo™ Cytotoxicity Assay Kit (Promega, Cat. No: J2381) according to manufacturer’s guidelines.

### Immunofluorescence (IF) and live cell imaging

To detect protein localization by immunofluorescence in fixed cells, cells were seeded on High Performance No.1.5 18 × 18 mm glass coverslips (Zeiss™ Cat. No: 474030-9000-000), and fixed with 4% PFA for 15 min, followed by permeabilization with 0.5% Triton X-100 for 5 min. Then, cells were blocked with 1.5% BSA for 1 hour at room temperature. Primary antibodies (ASC 1:200, AdipoGen: AG25B0006C100; LC3B 1:20, Santa Cruz Biotechnology: sc-271625) were diluted in 1.5% BSA and incubated overnight with fixed cells at 4 °C. After washing with 1 × DPBS 3 times, fluorescent secondary antibodies were 1: 1,000 diluted in 1.5% BSA and incubated for 1 hour at room temperature. Samples were mounted in VECTASHIELD antifade mounting medium (Vector Laboratories, Cat. No: H-1000-10) and sealed with Nail polish (Fisher Scientific, Cat. No: NC1849418).

For visualization of mScarlet-NLRP3, TGN38-mNeonGreen and SiR-tubulin in live cells, cells were differentiated and grown in glass bottom dishes (MatTek corporation, Cat. No: P35G-1.5–14-C) for 48 hours. Then cells were first washed once with PBS and the medium was replaced by RPMI 1640 medium with no phenol red (Thermo Fisher Scientific, Cat. No: 11835030) supplemented with 10% FBS, and placed back in the incubator for at least 1 hour. 1 μM SiR-Tubulin was used for 1-2 hours to stain the microtubule network in live cells. Live cell images were obtained at 37 °C with 5% CO_2_ condition.

All images were collected with oil-immersion 60X (1.4 numerical aperture) lenses on a spinning disk confocal on an inverted Nikon Ti fluorescence microscope, which was equipped with Nikon Perfect Focus System and Yokogawa CSU-X1 Confocal Scanner, and operated with NIS-Elements image acquisition software.

### Fluorescence recovery after photobleaching (FRAP) microscopy

mScarlet-NLRP3, or together with mNeonGreen-ASC reconstituted NLRP3 KO THP-1 cells were cultured on 35 mm glass-bottomed dishes (MatTek). FRAP microscopy for cytosolic or dTGN-localized NLRP3 before activation and dispersed NLRP3 after 10 min nigericin treatment was performed on a Deltavision OMX Blaze 3D Structured Illumination Microscope (SIM), equipped with sCMOS Cameras, and six laser beams (405, 445, 488, 514, 568 and 642 nm; 100 mW). To obtain optimal images, immersion oil with refractive indices of 1.516 was used. FRAP microscopy for NLRP3 and ASC specks after 30 min nigericin treatment was performed on a Leica STELLARIS 8 FALCON FLIM Microscope equipped with a 60 × 1.4 NA HC PL APO CS2 oil immersion objective and operated with the LASX acquisition software. The region of interest was photobleached and the recovery of fluorescence intensity within the region of interest was obtained for each experiment. Photobleaching and subsequent image acquisition were performed with 3 prebleached images and a sequence of 120 post-bleach images at 1 second intervals. Intensity recovery curves were normalized against photofading outside the bleaching area, calculated as percentage of recovery (98, 99), and fitted with a one phase exponential association curve by GraphPad Prism.

### Quantification and statistical analysis

The data used in this study were presented as mean ± standard deviation (SD) in triplicate experiments unless otherwise stated. Statistical analyses (two-tailed Student’s t-test, Mann-Whitney Test, linear regression and so on) were performed using existing software (GraphPad Prism 9 or 10). P < 0.05 was considered significant. Representative microscopy images were obtained from at least six independent cells. For the statistical significance and sample size of all graphs, please see figure legends.

---

**Fig. S1.**
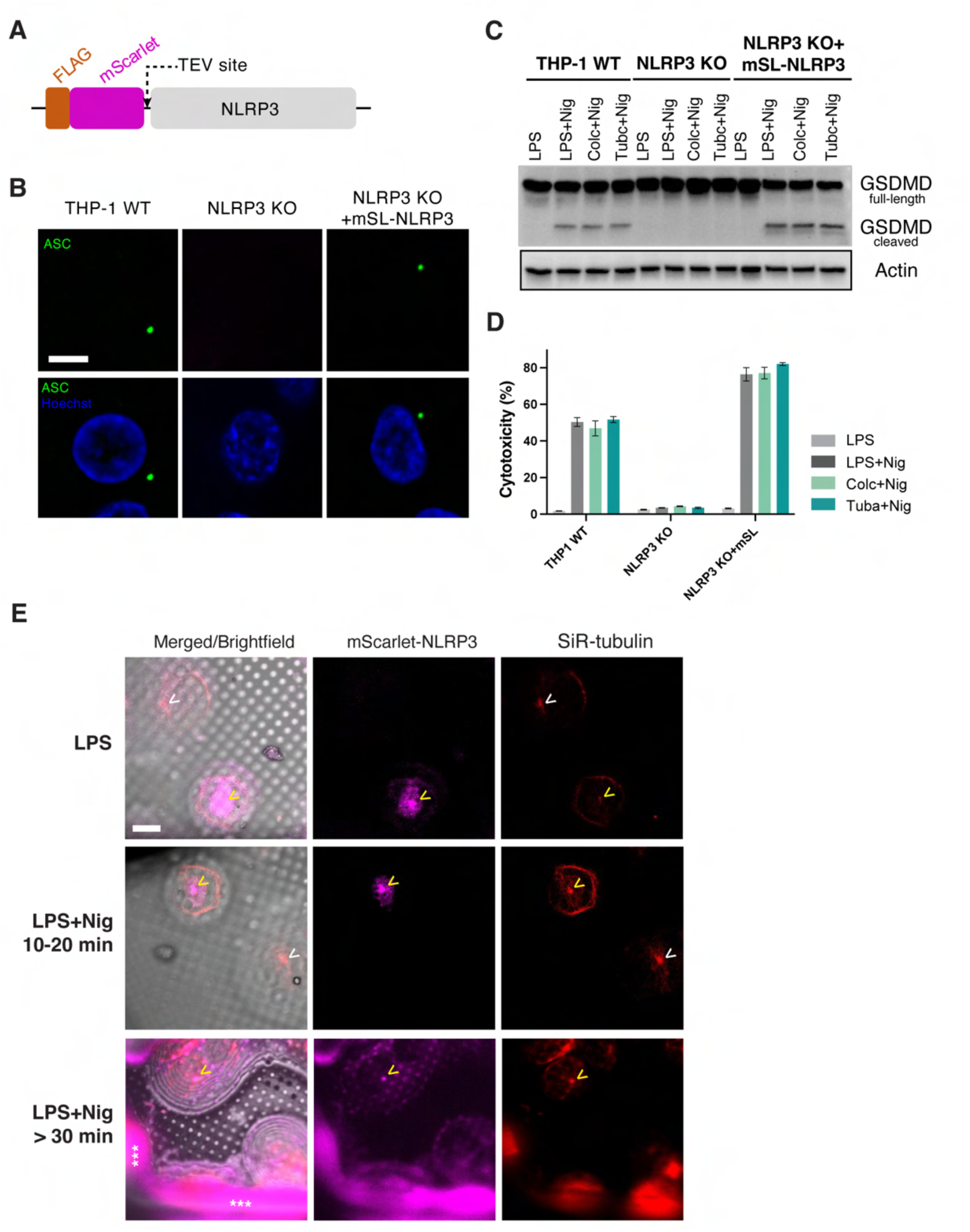
NLRP3 inflammasome activation leads to formation of a single punctum at the MTOC independent of retrograde transport in mScarlet-NLRP3 THP-1 cells. **(A)** A Schematic of the FLAG-mScarlet-NLRP3 construct. **(B)** Confocal imaging of ASC speck formation in THP-1 cells (WT, NLRP3 KO, and NLRP3 KO reconstituted with mSL-NLRP3) by IF. Cells were primed with1 μg/ml LPS for 4 hour and treated with 20 μM nigericin for 30 min. Green: ASC, Blue: nucleus (Scale bar=5 µm). **(C-D)** Western blotting (C) and LDH release assay (D) for NLRP3 activation in THP-1 cells (WT, NLRP3 KO, and NLRP3 KO reconstituted with mSL-NLRP3). LPS-primed cells were treated with 10 μM microtubule polymerization inhibitor Colchicine (Colc) or 20 μM HDAC6 inhibitor Tubacin (Tubc) for 2 hours before stimulation with 20 μM nigericin for 30 min. **(E)** Cryogenic fluorescent imaging of mSL-NLRP3 THP-1 cells before and after NLRP3 inflammasome activation. Cells were primed and activated following the same protocol in (A). NLRP3 inflammasome puncta are indicated as yellow arrows, and MTOC puncta in inactivated cells are indicated as white arrows. Grid bars are marked as asterisks (Scale bar=10 µm).

**Fig. S2.**
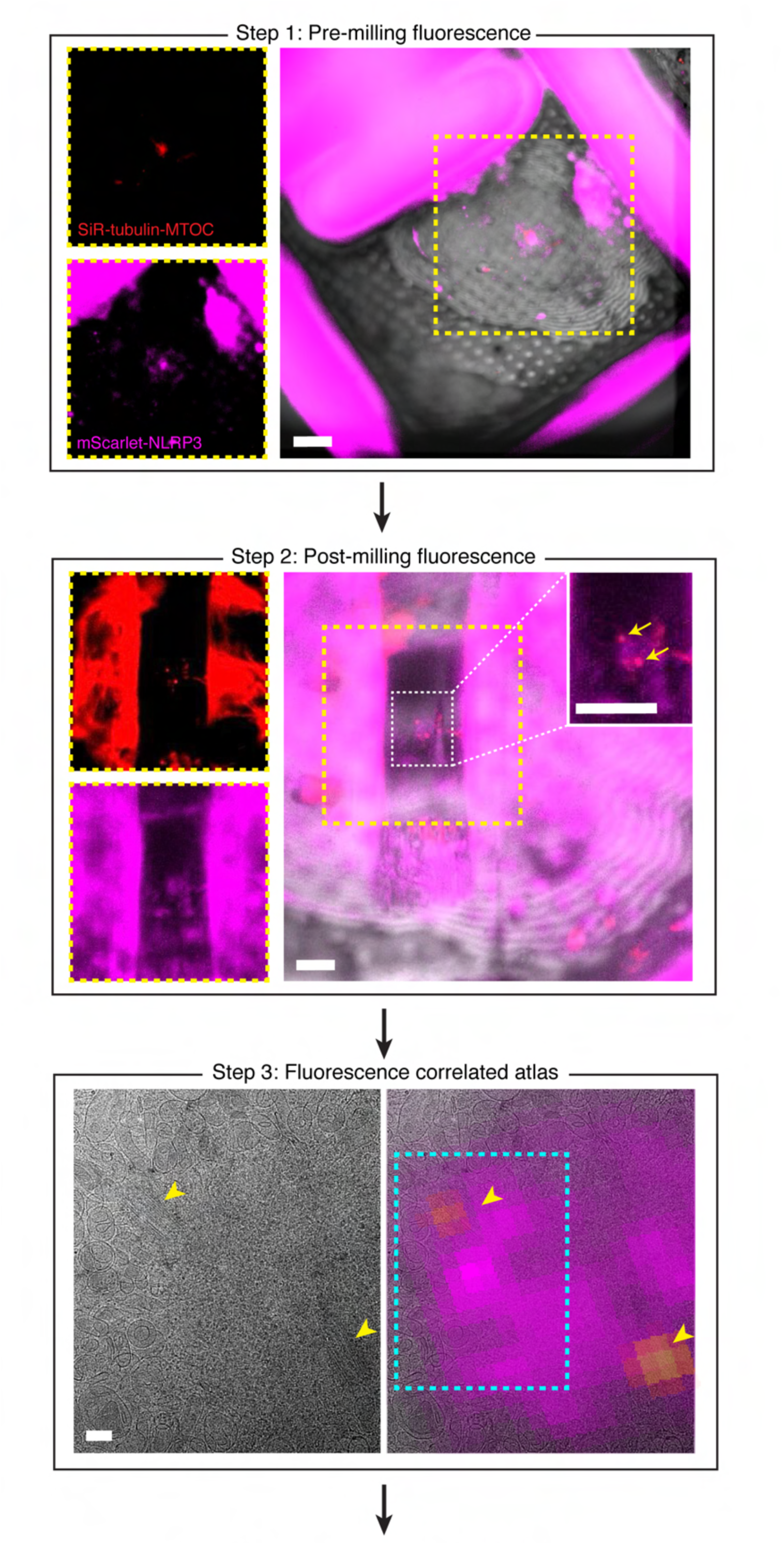
Fluorescence guided cryo-FIB-SEM milling workflow for targeting the NLRP3 inflammasome. Fluorescent images of stepwise targeted milling. The representative example was treated with LPS+Nig 10min. The same example was also shown in Fig. 1 C and Fig. 2D, I. Centrioles are indicated as yellow arrows. Imaging frame size is marked in cyan. Scale bar=10 µm for all panels.

**Fig. S3.**
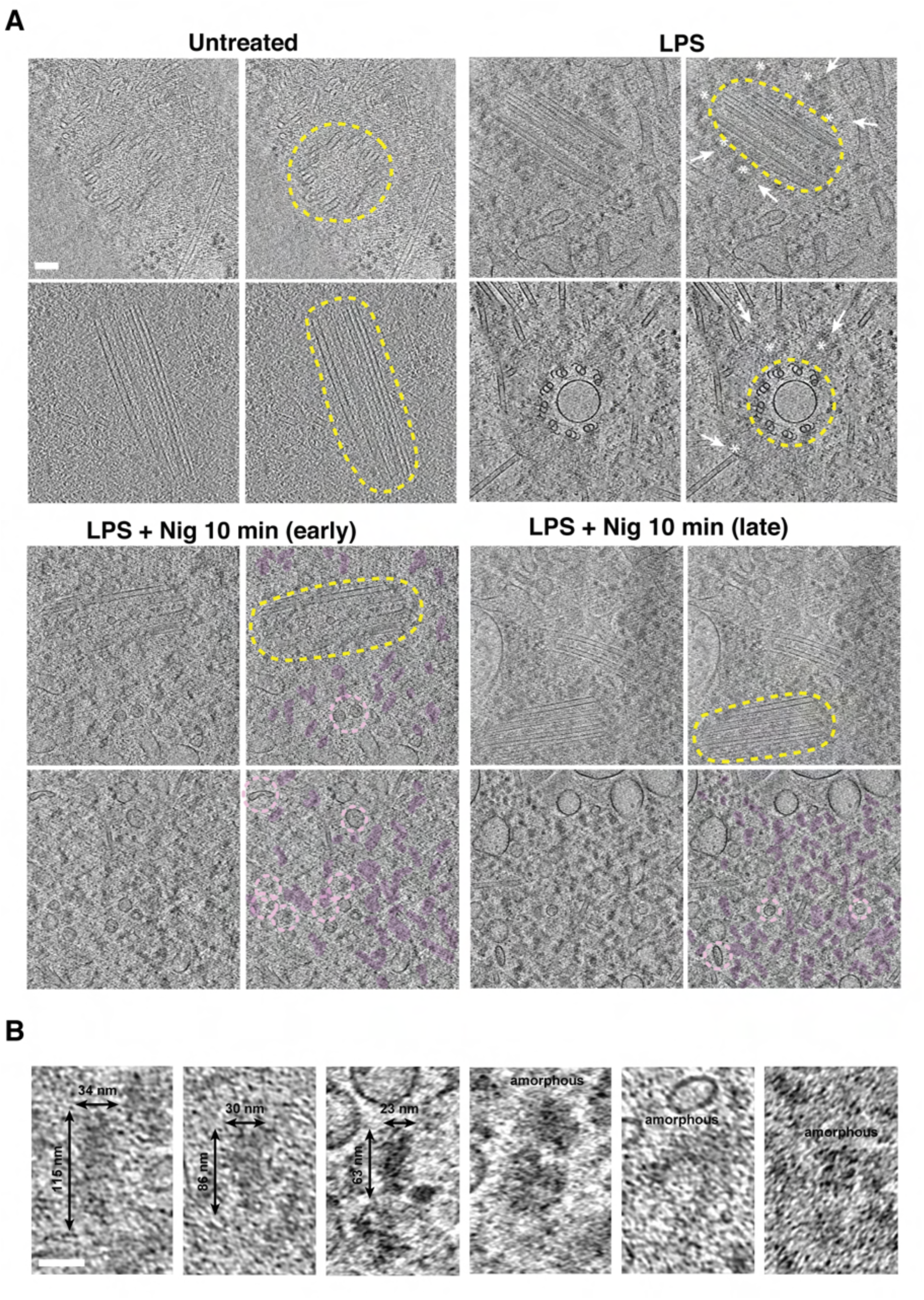
NLRP3 inflammasome forms condensate with pericentriolar material (PCM) and subject for autophagy. **(A)** Representative tomographic slices showing NLRP3 condensate in chronological order (scale bar=100 nm). Cellular features are marked as following: MTOC, yellow; NLRP3 condensate, magenta; NLRP3-associated vesicles, light pink. **(B)** Representative examples of NLRP3 condensate (scale bar=50 nm).

**Fig. S4.**
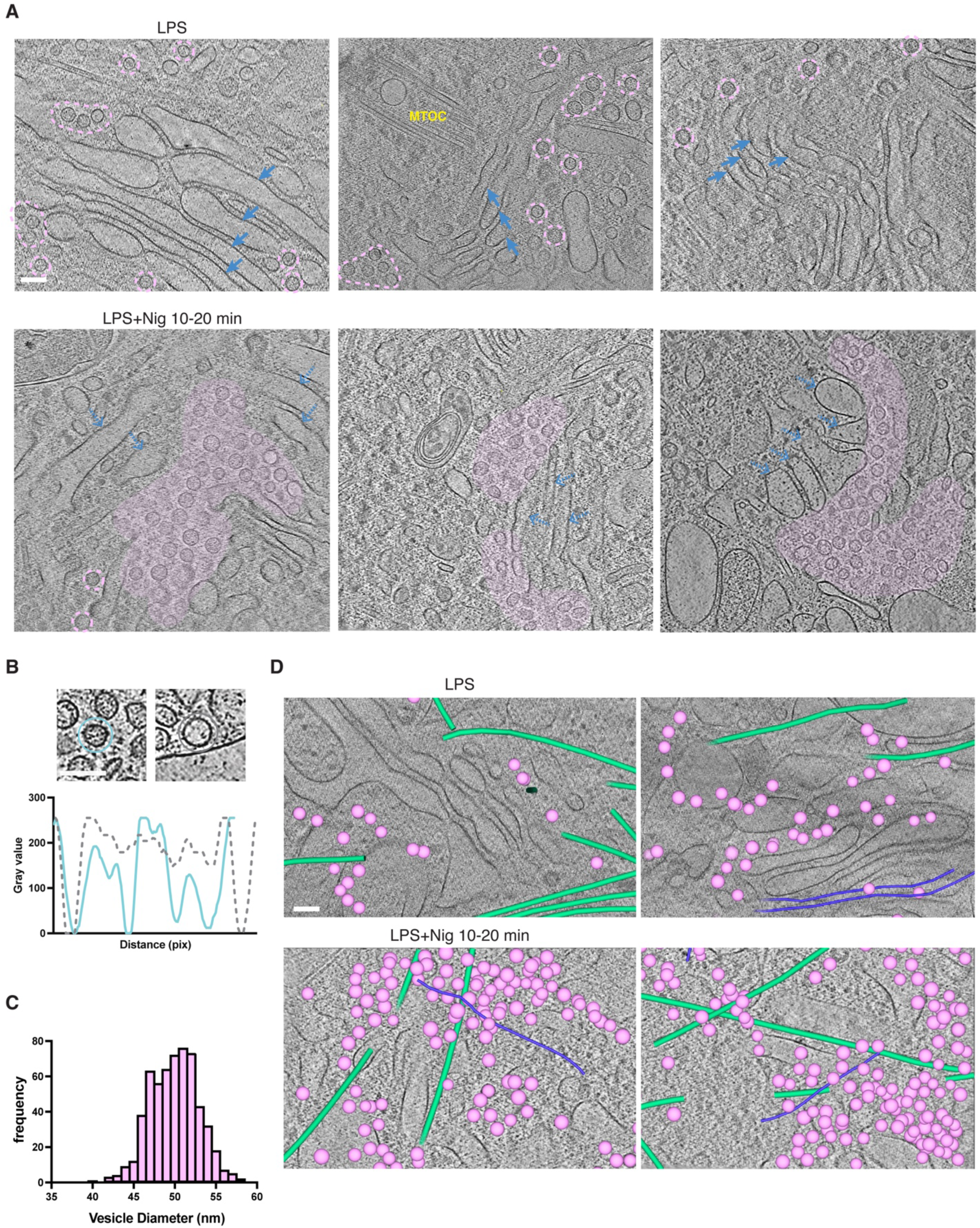
NLRP3-associated vesicles are trafficked to the MTOC in response to TGN dispersion via microtubule independent mechanism. **(A)** Representative examples of NLRP3-associated vesicles on TGN in primed cells and in the cytosol after addition of nigericin. Complete TGN cisternae are indicated as blue arrows and partial TGN structures or debris are indicated as dashed blue arrows. NLRP3-associated vesicles are circled or shaded in light pink (scale bar=100 nm). **(B)** NLRP3 associated vesicles are filled with protein densities compared to empty vesicles naturally present in cells (scale bar=50nm). Intensity profile of NLRP3-associated vesicles, blue line; empty vesicle, grey dashed line. **(C)** Diameter distribution of NLRP3-associated vesicles. We quantified 6 tomogram reconstructions collected after addition of nigericin for 10 min with a total of 578 vesicles. Each tomogram represents one cell obtained from different passage numbers during sample preparation. **(D)** Trafficking of NLRP3 associated vesicles is independent from retrograde transport. Cellular features are colored as following: NLRP3-associated vesicles, light pink; microtubules, green; intermediate filaments, purple (scale bar=100nm).

**Fig. S5.**
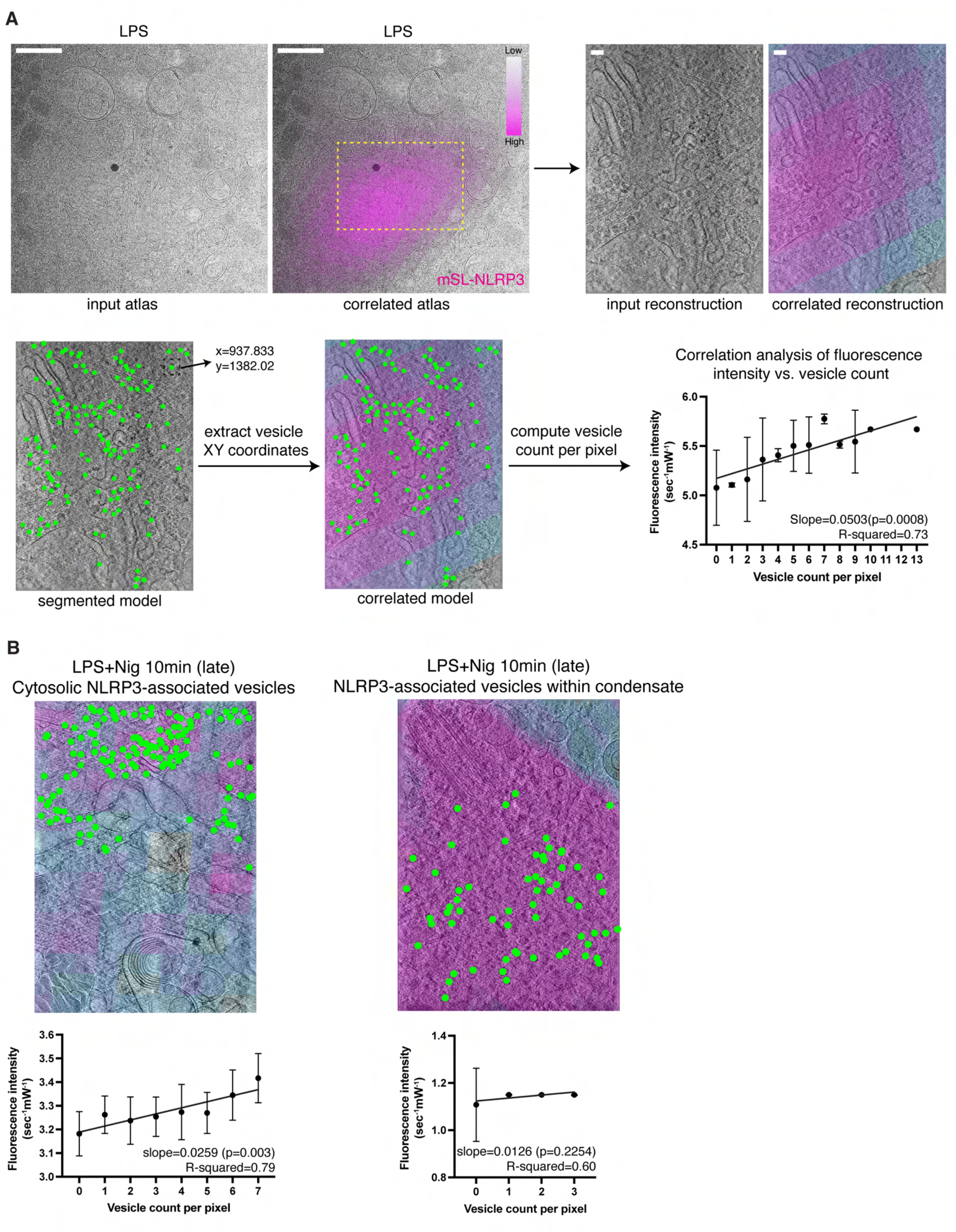
Fluorescence correlation analysis on NLRP3-associated vesicles. **(A)** Workflow of computing correlation between mScarlet fluorescence intensity and NLRP3-associated vesicle density (scale bar=1 µm for atlas; scale bar=100 nm for reconstructions). Fluorescence intensity has been normalized by exposure time and laser power. **(B)** Correlation analysis on NLRP3-associated vesicles observed in the cytosol and at the MTOC after 10 min-nigericin treatment.

**Fig. S6.**
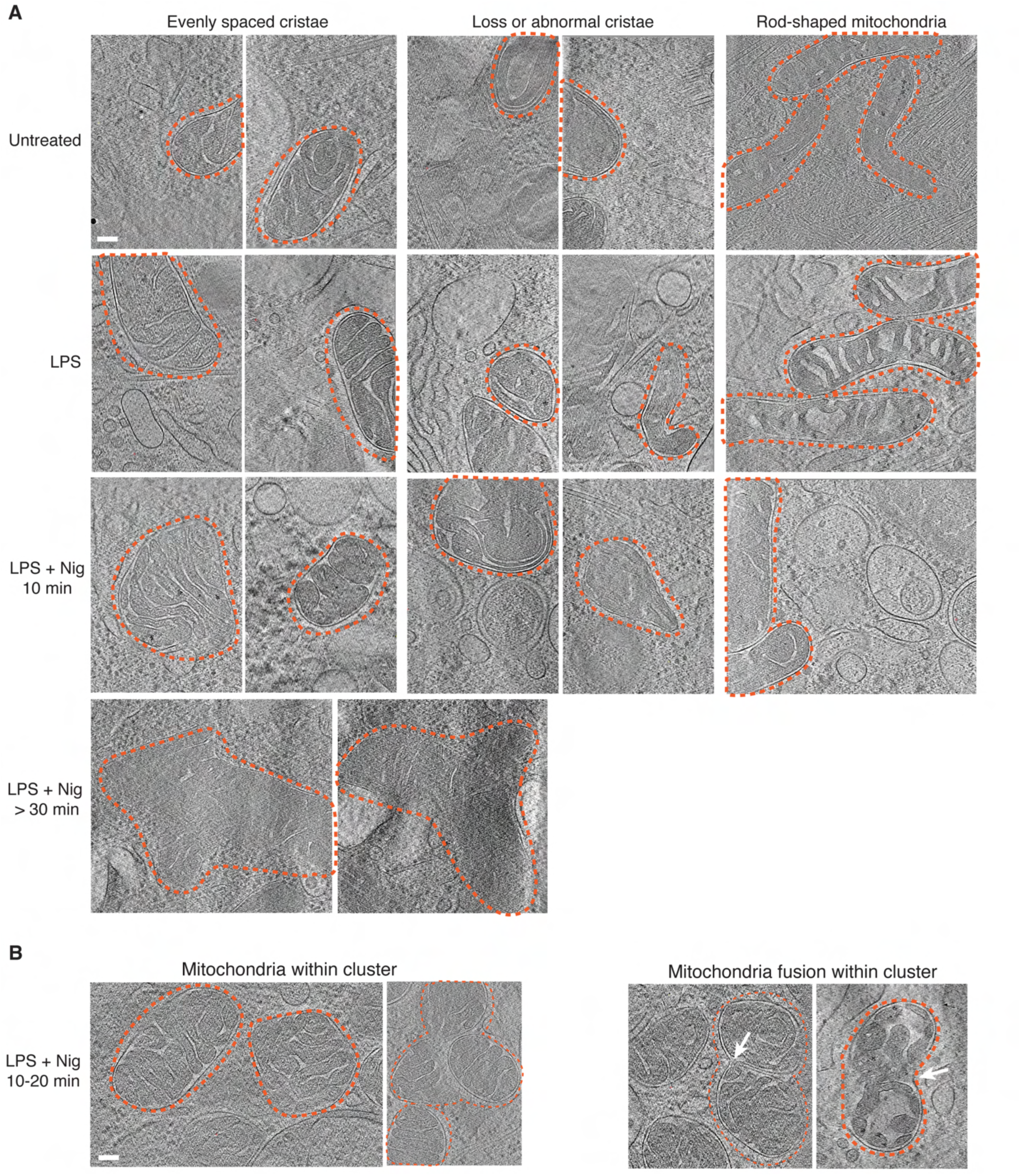
Mitochondria display a variety of morphology in response to NLRP3 inflammasome activation. **(A)** Gallery showing mitochondrial morphology before and after NLRP3 inflammasome activation. Mitochondria are circled in orange (scale bar=100 nm). **(B)** Representative examples of mitochondria observed within clusters after 10 min-nigericin treatment (scale bar=100 nm).

**Fig. S7.**
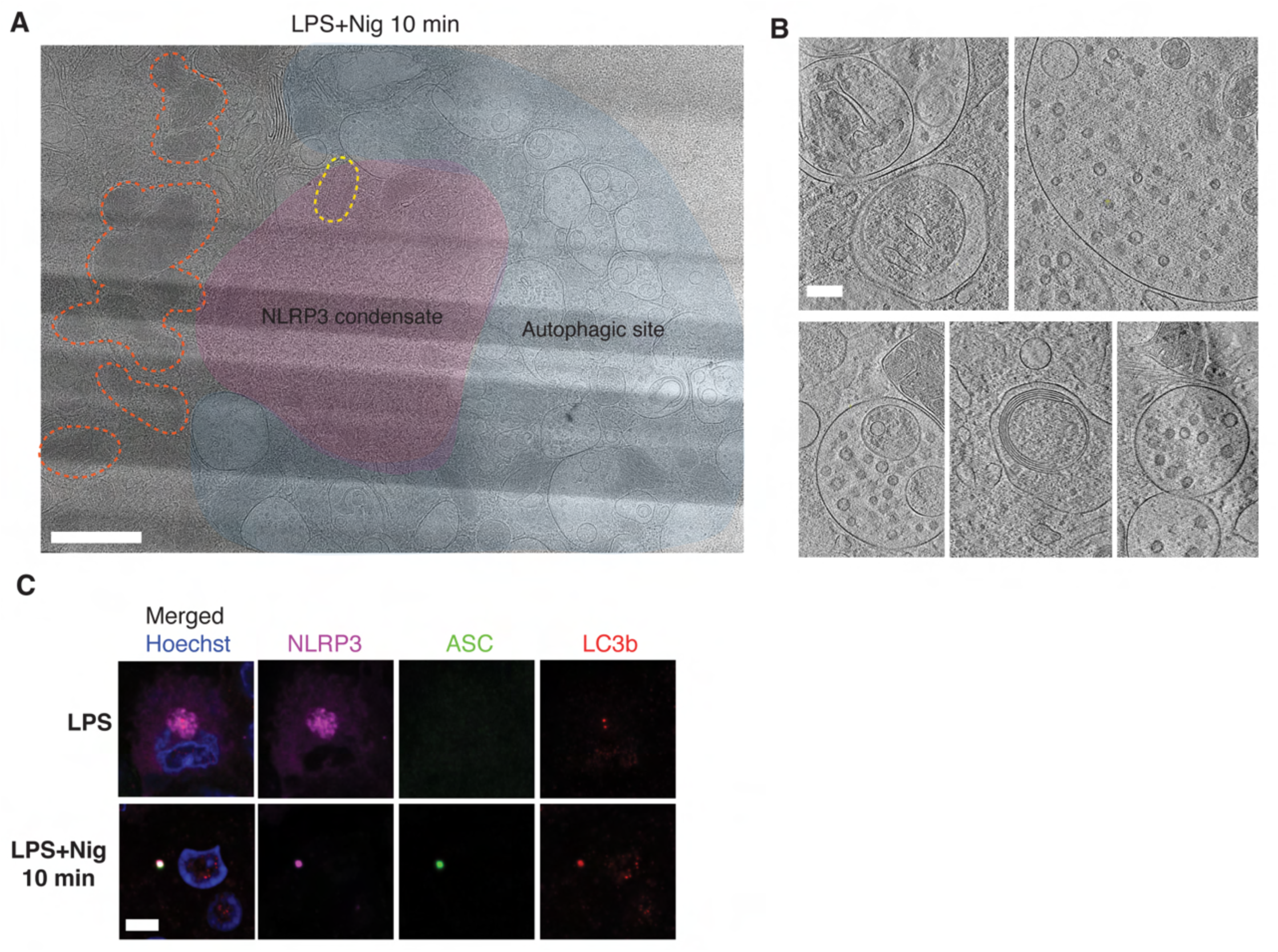
NLRP3 inflammasome condensate forms close contact with autophagic membranes. **(A)** Representative atlas showing NLRP3 condensate, autophagic membranes, and mitochondria in proximity. Cellular features are marked as following: NLRP3 condensate; magenta, autophagic membranes, blue; mitochondria, orange (scale bar=1 µm). **(B)** Gallery of autophagic membranes present in close contact with NLRP3 condensate (scale bar=100 nm). **(C)** Immunofluorescent imaging of NLRP3, ASC and LC3B in fixed cells. NT: none-treated; LPS: 1 μg/ml for 4 hours; LPS+Nig: LPS plus 20 μM nigericin for 30 min. Scale bar was shown.

## Notes

### Competing Interest Statement

HW is a co-founder and chair of scientific advisory board of Ventus Therapeutics. The remaining authors declare no competing interests.

## References and Notes

1. J. Fu, H. Wu, Structural Mechanisms of NLRP3 Inflammasome Assembly and Activation. Annu Rev Immunol 41, 301–316 (2023).

2. K. V. Swanson, M. Deng, J. P. Ting, The NLRP3 inflammasome: molecular activation and regulation to therapeutics. Nat Rev Immunol 19, 477–489 (2019).

3. K. Schroder, J. Tschopp, The inflammasomes. Cell 140, 821–832 (2010).

4. M. M. Gaidt, V. Hornung, The NLRP3 Inflammasome Renders Cell Death Pro-inflammatory. J Mol Biol 430, 133–141 (2018).

5. P. Broz, V. M. Dixit, Inflammasomes: mechanism of assembly, regulation and signalling. Nat Rev Immunol 16, 407–420 (2016).

6. X. Que, S. Zheng, Q. Song, H. Pei, P. Zhang, Fantastic voyage: The journey of NLRP3 inflammasome activation. Genes Dis 11, 819–829 (2024).

7. Z. Wang et al., NLRP3 Inflammasome and Inflammatory Diseases. Oxid Med Cell Longev 2020, 4063562 (2020).

8. A. Zahid, B. Li, A. J. K. Kombe, T. Jin, J. Tao, Pharmacological Inhibitors of the NLRP3 Inflammasome. Front Immunol 10, 2538 (2019).

9. D. Wu et al., Inflammasome Meets Centrosome: Understanding the Emerging Role of Centrosome in Controlling Inflammasome Activation. Front Immunol 13, 826106 (2022).

10. L. Xiao, V. G. Magupalli, H. Wu, Cryo-EM structures of the active NLRP3 inflammasome disc. Nature 613, 595–600 (2023).

11. Y. Dong et al., Structural transitions enable interleukin-18 maturation and signaling. Immunity, (2024).

12. J. Lieberman, H. Wu, J. C. Kagan, Gasdermin D activity in inflammation and host defense. Sci Immunol 4, (2019).

13. V. Hornung et al., AIM2 recognizes cytosolic dsDNA and forms a caspase-1-activating inflammasome with ASC. Nature 458, 514–518 (2009).

14. V. G. Magupalli et al., HDAC6 mediates an aggresome-like mechanism for NLRP3 and pyrin inflammasome activation. Science 369, (2020).

15. J. Wu, T. Fernandes-Alnemri, E. S. Alnemri, Involvement of the AIM2, NLRC4, and NLRP3 inflammasomes in caspase-1 activation by Listeria monocytogenes. J Clin Immunol 30, 693–702 (2010).

16. X. Li et al., MARK4 regulates NLRP3 positioning and inflammasome activation through a microtubule-dependent mechanism. Nat Commun 8, 15986 (2017).

17. H. Sharif et al., Structural mechanism for NEK7-licensed activation of NLRP3 inflammasome. Nature 570, 338–343 (2019).

18. X. Yu et al., Structural basis for the oligomerization-facilitated NLRP3 activation. Nature communications 15, 1164 (2024).

19. L. Andreeva et al., NLRP3 cages revealed by full-length mouse NLRP3 structure control pathway activation. Cell 184, 6299–6312.e6222 (2021).

20. I. V. Hochheiser et al., Structure of the NLRP3 decamer bound to the cytokine release inhibitor CRID3. Nature 604, 184–189 (2022).

21. U. Ohto et al., Structural basis for the oligomerization-mediated regulation of NLRP3 inflammasome activation. Proc Natl Acad Sci U S A 119, e2121353119 (2022).

22. A. Stutz, G. L. Horvath, B. G. Monks, E. Latz, ASC speck formation as a readout for inflammasome activation. Methods Mol Biol 1040, 91–101 (2013).

23. Y. Liu et al., Cryo-electron tomography of NLRP3-activated ASC complexes reveals organelle co-localization. Nature communications 14, 7246 (2023).

24. Y. Li et al., Cryo-EM structures of ASC and NLRC4 CARD filaments reveal a unified mechanism of nucleation and activation of caspase-1. Proc Natl Acad Sci U S A 115, 10845–10852 (2018).

25. A. Lu et al., Unified polymerization mechanism for the assembly of ASC-dependent inflammasomes. Cell 156, 1193–1206 (2014).

26. C. M. Hampton et al., Correlated fluorescence microscopy and cryo-electron tomography of virus-infected or transfected mammalian cells. Nat Protoc 12, 150–167 (2017).

27. G. H. Wu et al., Multi-scale 3D Cryo-Correlative Microscopy for Vitrified Cells. Structure 28, 1231–1237.e1233 (2020).

28. S. D. Carter, J. I. Mamede, T. J. Hope, G. J. Jensen, Correlated cryogenic fluorescence microscopy and electron cryo-tomography shows that exogenous TRIM5α can form hexagonal lattices or autophagy aggregates in vivo. Proc Natl Acad Sci U S A 117, 29702–29711 (2020).

29. D. B. Boltje et al., A cryogenic, coincident fluorescence, electron, and ion beam microscope. Elife 11, (2022).

30. M. Lamkanfi, V. M. Dixit, Mechanisms and functions of inflammasomes. Cell 157, 1013–1022 (2014).

31. R. Muñoz-Planillo et al., K⁺ efflux is the common trigger of NLRP3 inflammasome activation by bacterial toxins and particulate matter. Immunity 38, 1142–1153 (2013).

32. G. Lukinavicius et al., Fluorogenic probes for live-cell imaging of the cytoskeleton. Nat Methods 11, 731–733 (2014).

33. X. Zhang et al., Molecular mechanisms of stress-induced reactivation in mumps virus condensates. Cell 186, 1877–1894.e1827 (2023).

34. Q. Guo et al., In Situ Structure of Neuronal C9orf72 Poly-GA Aggregates Reveals Proteasome Recruitment. Cell 172, 696–705.e612 (2018).

35. M. H. Laporte et al., Time-series reconstruction of the molecular architecture of human centriole assembly. Cell 187, 2158–2174.e2119 (2024).

36. M. LeGuennec, N. Klena, G. Aeschlimann, V. Hamel, P. Guichard, Overview of the centriole architecture. Curr Opin Struct Biol 66, 58–65 (2021).

37. M. Paintrand, M. Moudjou, H. Delacroix, M. Bornens, Centrosome organization and centriole architecture: their sensitivity to divalent cations. J Struct Biol 108, 107–128 (1992).

38. S. Li et al., ELI trifocal microscope: a precise system to prepare target cryo-lamellae for in situ cryo-ET study. Nat Methods 20, 276–283 (2023).

39. H. Fujita, Y. Yoshino, N. Chiba, Regulation of the centrosome cycle. Mol Cell Oncol 3, e1075643 (2016).

40. X. Jiang et al., Condensation of pericentrin proteins in human cells illuminates phase separation in centrosome assembly. J Cell Sci 134, (2021).

41. J. B. Woodruff et al., The Centrosome Is a Selective Condensate that Nucleates Microtubules by Concentrating Tubulin. Cell 169, 1066–1077.e1010 (2017).

42. T. Consolati et al., Microtubule Nucleation Properties of Single Human γTuRCs Explained by Their Cryo-EM Structure. Dev Cell 53, 603–617.e608 (2020).

43. V. Guillet et al., Crystal structure of γ-tubulin complex protein GCP4 provides insight into microtubule nucleation. Nat Struct Mol Biol 18, 915–919 (2011).

44. J. M. Kollman, A. Merdes, L. Mourey, D. A. Agard, Microtubule nucleation by γ-tubulin complexes. Nat Rev Mol Cell Biol 12, 709–721 (2011).

45. A. Vertii et al., The Centrosome Undergoes Plk1-Independent Interphase Maturation during Inflammation and Mediates Cytokine Release. Dev Cell 37, 377–386 (2016).

46. M. T. Bertran et al., Nek9 is a Plk1-activated kinase that controls early centrosome separation through Nek6/7 and Eg5. EMBO J 30, 2634–2647 (2011).

47. E. Smith et al., Differential control of Eg5-dependent centrosome separation by Plk1 and Cdk1. EMBO J 30, 2233–2245 (2011).

48. E. Vitiello et al., Acto-myosin force organization modulates centriole separation and PLK4 recruitment to ensure centriole fidelity. Nat Commun 10, 52 (2019).

49. M. S. Ladinsky, D. N. Mastronarde, J. R. McIntosh, K. E. Howell, L. A. Staehelin, Golgi structure in three dimensions: functional insights from the normal rat kidney cell. J Cell Biol 144, 1135–1149 (1999).

50. M. H. Dunlop et al., Land-locked mammalian Golgi reveals cargo transport between stable cisternae. Nat Commun 8, 432 (2017).

51. D. M. Williams, A. A. Peden, S-acylation of NLRP3 provides a nigericin sensitive gating mechanism that controls access to the Golgi. bioRxiv, 2023.2011.2014.566891 (2023).

52. N. A. Schmacke et al., IKKβ primes inflammasome formation by recruiting NLRP3 to the trans-Golgi network. Immunity 55, 2271–2284.e2277 (2022).

53. J. Chen, Z. J. Chen, PtdIns4P on dispersed trans-Golgi network mediates NLRP3 inflammasome activation. Nature 564, 71–76 (2018).

54. G. dos Santos et al., Vimentin regulates activation of the NLRP3 inflammasome. Nature communications 6, 6574 (2015).

55. T. Lang et al., Macrophage migration inhibitory factor is required for NLRP3 inflammasome activation. Nature communications 9, 2223 (2018).

56. P. Gurung, J. R. Lukens, T. D. Kanneganti, Mitochondria: diversity in the regulation of the NLRP3 inflammasome. Trends Mol Med 21, 193–201 (2015).

57. Q. Liu, D. Zhang, D. Hu, X. Zhou, Y. Zhou, The role of mitochondria in NLRP3 inflammasome activation. Mol Immunol 103, 115–124 (2018).

58. M. Yabal, D. J. Calleja, D. S. Simpson, K. E. Lawlor, Stressing out the mitochondria: Mechanistic insights into NLRP3 inflammasome activation. J Leukoc Biol 105, 377–399 (2019).

59. R. Zhou, A. S. Yazdi, P. Menu, J. Tschopp, A role for mitochondria in NLRP3 inflammasome activation. Nature 469, 221–225 (2011).

60. A. Kaasik, D. Safiulina, A. Zharkovsky, V. Veksler, Regulation of mitochondrial matrix volume. Am J Physiol Cell Physiol 292, C157–163 (2007).

61. P. Huang, T. Yu, Y. Yoon, Mitochondrial clustering induced by overexpression of the mitochondrial fusion protein Mfn2 causes mitochondrial dysfunction and cell death. Eur J Cell Biol 86, 289–302 (2007).

62. Y. Huo, W. Sun, T. Shi, S. Gao, M. Zhuang, The MFN1 and MFN2 mitofusins promote clustering between mitochondria and peroxisomes. Commun Biol 5, 423 (2022).

63. S. Agarwal, S. Ganesh, Perinuclear mitochondrial clustering, increased ROS levels, and HIF1 are required for the activation of HSF1 by heat stress. J Cell Sci 133, (2020).

64. A. B. Al-Mehdi et al., Perinuclear mitochondrial clustering creates an oxidant-rich nuclear domain required for hypoxia-induced transcription. Sci Signal 5, ra47 (2012).

65. G. Du et al., ROS-dependent S-palmitoylation activates cleaved and intact gasdermin D. Nature 630, 437–446 (2024).

66. R. Miao et al., Gasdermin D permeabilization of mitochondrial inner and outer membranes accelerates and enhances pyroptosis. Immunity 56, 2523–2541.e2528 (2023).

67. H. Shi et al., NLRP3 activation and mitosis are mutually exclusive events coordinated by NEK7, a new inflammasome component. Nat Immunol 17, 250–258 (2016).

68. S. Bai, F. Martin-Sanchez, D. Brough, G. Lopez-Castejon, Pyroptosis leads to loss of centrosomal integrity in macrophages. bioRxiv, 2023.2011.2022.568260 (2023).

69. S. Alberti, A. A. Hyman, Biomolecular condensates at the nexus of cellular stress, protein aggregation disease and ageing. Nat Rev Mol Cell Biol 22, 196–213 (2021).

70. D. M. Shapiro, M. Ney, S. A. Eghtesadi, A. Chilkoti, Protein Phase Separation Arising from Intrinsic Disorder: First-Principles to Bespoke Applications. J Phys Chem B 125, 6740–6759 (2021).

71. F. Garcia Quiroz et al., Intrinsically disordered proteins access a range of hysteretic phase separation behaviors. Sci Adv 5, eaax5177 (2019).

72. M. Du, Z. J. Chen, DNA-induced liquid phase condensation of cGAS activates innate immune signaling. Science 361, 704–709 (2018).

73. F. Jobe, J. Simpson, P. Hawes, E. Guzman, D. Bailey, Respiratory Syncytial Virus Sequesters NF-κB Subunit p65 to Cytoplasmic Inclusion Bodies To Inhibit Innate Immune Signaling. J Virol 94, (2020).

74. C. Shen et al., Phase separation drives RNA virus-induced activation of the NLRP6 inflammasome. Cell 184, 5759–5774.e5720 (2021).

75. X. Su et al., Phase separation of signaling molecules promotes T cell receptor signal transduction. Science 352, 595–599 (2016).

76. X. Yu et al., The STING phase-separator suppresses innate immune signalling. Nat Cell Biol 23, 330–340 (2021).

77. H. H. Park et al., Death domain assembly mechanism revealed by crystal structure of the oligomeric PIDDosome core complex. Cell 128, 533–546 (2007).

78. S. C. Lin, Y. C. Lo, H. Wu, Helical assembly in the MyD88-IRAK4-IRAK2 complex in TLR/IL-1R signalling. Nature 465, 885–890 (2010).

79. H. Wu, Higher-order assemblies in a new paradigm of signal transduction. Cell 153, 287–292 (2013).

80. H. Wu, M. Fuxreiter, The Structure and Dynamics of Higher-Order Assemblies: Amyloids, Signalosomes, and Granules. Cell 165, 1055–1066 (2016).

81. L. Cui et al., Vesicle trafficking and vesicle fusion: mechanisms, biological functions, and their implications for potential disease therapy. Mol Biomed 3, 29 (2022).

82. R. Fernández-Busnadiego et al., Quantitative analysis of the native presynaptic cytomatrix by cryoelectron tomography. J Cell Biol 188, 145–156 (2010).

83. J. C. Nelson, A. K. Stavoe, D. A. Colón-Ramos, The actin cytoskeleton in presynaptic assembly. Cell Adh Migr 7, 379–387 (2013).

84. H. Qiu et al., Short-distance vesicle transport via phase separation. Cell 187, 2175–2193.e2121 (2024).

85. D. Milovanovic, Y. Wu, X. Bian, P. De Camilli, A liquid phase of synapsin and lipid vesicles. Science 361, 604–607 (2018).

86. T. C. Südhof, P. De Camilli, H. Niemann, R. Jahn, Membrane fusion machinery: insights from synaptic proteins. Cell 75, 1–4 (1993).

87. F. Navone, P. Greengard, P. De Camilli, Synapsin I in nerve terminals: selective association with small synaptic vesicles. Science 226, 1209–1211 (1984).

88. G. Lukinavičius et al., Fluorogenic probes for live-cell imaging of the cytoskeleton. Nat Methods 11, 731–733 (2014).

89. V. Lam, E. Villa, Practical Approaches for Cryo-FIB Milling and Applications for Cellular Cryo-Electron Tomography. Methods Mol Biol 2215, 49–82 (2021).

90. F. R. Wagner et al., Preparing samples from whole cells using focused-ion-beam milling for cryo-electron tomography. Nat Protoc 15, 2041–2070 (2020).

91. D. N. Mastronarde, SerialEM: A Program for Automated Tilt Series Acquisition on Tecnai Microscopes Using Prediction of Specimen Position. Microscopy and Microanalysis 9, 1182–1183 (2003).

92. D. N. Mastronarde, Automated electron microscope tomography using robust prediction of specimen movements. J Struct Biol 152, 36–51 (2005).

93. D. Tegunov, P. Cramer, Real-time cryo-electron microscopy data preprocessing with Warp. Nat Methods 16, 1146–1152 (2019).

94. D. Tegunov, L. Xue, C. Dienemann, P. Cramer, J. Mahamid, Multi-particle cryo-EM refinement with M visualizes ribosome-antibiotic complex at 3.5 Å in cells. Nat Methods 18, 186–193 (2021).

95. J. R. Kremer, D. N. Mastronarde, J. R. McIntosh, Computer visualization of three-dimensional image data using IMOD. J Struct Biol 116, 71–76 (1996).

96. D. N. Mastronarde, S. R. Held, Automated tilt series alignment and tomographic reconstruction in IMOD. J Struct Biol 197, 102–113 (2017).

97. E. C. Meng et al., UCSF ChimeraX: Tools for structure building and analysis. Protein Sci 32, e4792 (2023).

98. M. Dundr et al., A kinetic framework for a mammalian RNA polymerase in vivo. Science 298, 1623–1626 (2002).

99. R. D. Phair, S. A. Gorski, T. Misteli, Measurement of dynamic protein binding to chromatin in vivo, using photobleaching microscopy. Methods Enzymol 375, 393–414 (2004).

